# AMPAR immunization induces progressive autoimmune encephalitis with autoreactive B cells in the brain

**DOI:** 10.64898/2026.05.26.727986

**Authors:** Justus B. H. Wilke, George Celis, Neo Yixuan Peng, Natalie Sheldon, Cathy J. Spangler, Dushyant K. Srivastava, April Goehring, Harald Prüss, Gary Westbrook, Lauren B. Rodda, Eric Gouaux

## Abstract

AMPA and NMDA receptors are central to synaptic plasticity and cognitive function. In anti-NMDA receptor and anti-AMPA receptor (AMPAR) encephalitis, autoantibodies targeting these receptors disrupt synaptic signaling, leading to severe neuropsychiatric symptoms. However, the cellular autoimmune responses and source of pathogenic autoantibodies during onset and progression of central nervous system (CNS) pathology remain poorly understood. By immunizing mice with intact AMPARs in proteoliposomes, we developed a mouse model of anti-AMPAR encephalitis. Mice developed rapidly progressing neuropsychiatric deficits, autoantibodies targeting the AMPAR amino-terminal domain (ATD) and IgG deposition in the brain, accompanied by reduced AMPAR detection. Throughout disease onset and progression, AMPAR-ATD-specific non-proliferating plasma cells and plasmablasts accumulated in the brain and were predominantly localized in AMPAR-expressing brain parenchyma. In contrast, differentiated AMPAR-ATD specific B cells were far less enriched in peripheral lymphoid tissues. Our results suggest that humoral autoimmune responses directly in the CNS drives disease progression in anti-AMPAR encephalitis.

## INTRODUCTION

Autoimmune encephalitides (AEs) are brain disorders characterized by severe psychiatric and neurological symptoms as well as autoantibodies against specific central nervous system (CNS) proteins^1–4^. In anti-NMDAR encephalitis (NMDAR-AE) and anti-AMPAR encephalitis (AMPAR-AE), these autoantibodies target two closely related ionotropic glutamate receptors that are crucial to excitatory synaptic function^1^. First described in 2009, AMPAR-AE was discovered in ten adults presenting with subacute limbic encephalitis, characterized by brain inflammation and anti-AMPAR autoantibodies in cerebrospinal fluid^5^. Case series and systematic reviews subsequently revealed heterogeneous presentations, frequent tumor associations, and variable outcomes^5–11^. AMPAR-AE is thought to be caused by an autoimmune response against AMPAR in affected brain regions possibly initiated by AMPAR-expression in the non-CNS tumors in these patients^5^. However, the role and location of the immune response in AMPAR-AE symptom onset and progression remain poorly understood. Current treatments with immunosuppressive steroids, B cell depletion, and antibody-replacing plasmapheresis have some success, but relapses are common and functional deficits remain in approximately 80% of adult and 50% of pediatric cases^10^. Further mechanistic insights are needed to facilitate targeted therapies with durable benefits.

AMPARs are glutamate-activated, non-selective cation channels, typically composed of subunits GluA1-GluA4, that mediate fast excitatory neurotransmission and are central to the development and function of the nervous system^12,13^. Because AMPARs play a crucial role in excitatory synaptic transmission, it has been hypothesized that disease-associated autoantibodies mediate cognitive impairments. AMPAR autoantibodies in AMPAR-AE patients preferentially target the GluA2 and/or GluA1 subunits with binding sites primarily localized to the amino terminal domain (ATD)^5,7,14^ resulting in AMPAR internalization *in vitro*^5,14–16^. The *in vivo* pathogenicity of patient-derived GluA2 IgG has been demonstrated by passive autoantibody transfer into the brain ventricles and hippocampus of mice which reduced synaptic GluA2-containing AMPARs accompanied by alterations in miniature excitatory postsynaptic currents that resembled neuron-specific GluA2 knockout mice^15^. Furthermore, passive transfer of patient-derived GluA2-autoantibodies into the cerebrospinal fluid of mice resulted in progressive memory deficits that reversed within days after cessation of antibody infusion^15^, highlighting that continuous autoantibody generation by ASCs would be required for prolonged symptoms. In affected patients, AMPAR autoantibodies make up a higher proportion of IgG antibody in cerebrospinal fluid than in matched serum^5^, suggesting that ASCs in the CNS are the source of pathogenic AMPAR autoantibodies.

While passive autoantibody transfers have been instrumental in identifying antibody-mediated effects, they lack autoantigen-experienced T and B cells and do not fully recapitulate the CNS immunopathology and disease course of AE patients^17^. Hence, we developed an active immunization mouse model by immunizing mice with proteoliposomes containing homotetrameric GluA2 receptors. Immunization initiated a neuroinflammatory cascade with accumulation of T cells, B cells, and ASCs within AMPAR-expressing brain regions. To track and localize AMPAR-specific B lineage cells we developed specific AMPAR-ATD antigen probes and discovered that AMPAR-autoantibody synthesizing ASCs and AMPAR-specific MBCs accumulated within the meninges and brain parenchyma as the disease progressed. Our findings suggest that the humoral autoimmune response in AMPAR-AE is perpetuated in the brain and meninges. Our findings provide insights into disease mechanisms and highlight possible new targets for therapy.

## RESULTS

### Immunization with AMPAR-proteoliposomes induces AMPAR-AE-like disease in mice

To enable a deep and comprehensive understanding of AMPAR-AE and the roles of autoantibodies and neuroinflammation in AMPAR-AE, we developed an active immunization mouse model as well as new molecular tools. AMPAR-AE has been associated with peripheral tumors expressing AMPARs and most AMPAR-AE patients have autoantibodies targeting the AMPAR GluA2 subunit^5,7,14^. To induce an autoimmune response against the intact and folded receptor, we immunized mice intraperitoneally with proteoliposomes composed of homotetrameric GluA2-receptors embedded in liposomes (‘GluA2-PLP mice’). Control mice were immunized with non-protein containing liposomes and equivalent doses of adjuvant (‘Control’). To assess the clinical phenotype and disease course, mice were monitored daily for signs of encephalitis, as previously described for NMDAR-AE mice^18^. We assessed CNS histopathology and GluA2-specific autoimmunity in the CNS and lymphoid tissues at 3±1 days after the onset of hyperlocomotion (‘Onset’) and the remaining mice at day 56 post-primary immunization or earlier if they reached a humane endpoint (‘Progressed’) (Fig. 1A).

**Fig. 1.**
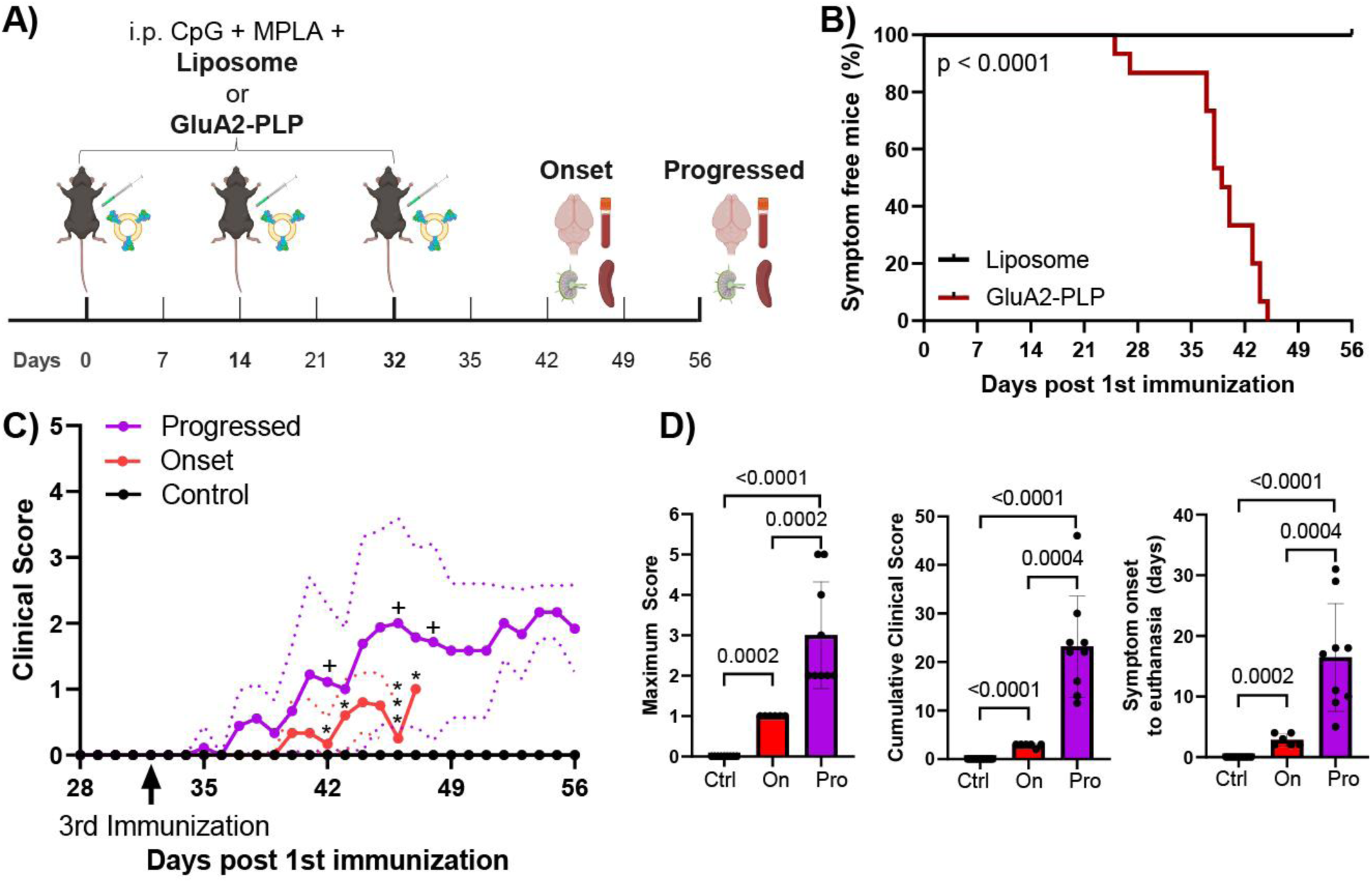
Timeline and clinical scoring of AMPAR-AE mouse model. (**A**) Immunizations and analyses of AMPAR-AE mice as a function of time (C57BL6 females, GluA2-PLP immunized n=15, liposome immunized n=15, Onset n=6 per group, Progressed n=9 per group; 2 experiments; created with BioRender). (**B**) Mice immunized with GluA2-PLP or liposomes were monitored for symptoms over 56 days (log-rank Mantel-Cox test). (**C**) Disease course as measured by clinical score over time for each group (mean (bold line) ± SD (dotted lines and error bars), ^+^humane endpoint reached, *mouse analyzed) and (**D**) disease severity in the experimental groups by clinical score and duration of symptoms (Ctrl, liposome Control; On, Onset; Pro, Progressed; Mann Whitney U tests).

Within 25-45 days of primary immunization, all GluA2-PLP mice developed pronounced hyperactivity characterized by excessive response to handling as well as spontaneous hyperlocomotion and erratic movements, particularly of the head (Fig. 1B, and Video S1). While we found substantial inter-individual differences, GluA2-PLP mice progressed towards more severe neurological symptoms with time after immunization (Fig. 1C and 1D). This included hyperactivity interrupted by episodes of behavioral quiescence, during which mice displayed body and head tremors (Video S2), as well as movement disorders, kyphosis and lethargy. Atypical movements started with light tottering (Video S3), which developed into severe ataxia and choreiform head movements (Video S4) reminiscent of stargazer mice, which lack synaptic GluA2-containing AMPAR in cerebellar granule cells^19,20^. Liposome immunized control mice did not display any symptoms or aberrant behaviors.

### Anti-AMPAR autoantibodies primarily target the GluA2 ATD and accumulate in the CNS

Anti-GluA2 IgG autoantibodies have been associated with pathological events in AMPAR-AE^5,15^. GluA2-PLP immunization induced anti-GluA2 IgG autoantibodies in mouse plasma as measured by live-cell based assay (Fig. 2A) and non-denaturing ELISA (Fig. S1A) coated with homotetrameric GluA2-receptors^21^. A well characterized, 3D epitope-binding monoclonal mouse anti-GluA2-ATD antibody (clone 15F1, Fig. 2B) was used as the standard^22–24^. GluA2-PLP mice had higher plasma concentrations of anti-GluA2 IgG than Control mice, which were below the detection limit. Despite the higher clinical score of Progressed relative to Onset mice, the plasma anti-GluA2 IgG concentration was similarly high (2.9-16.8μM) in both groups (Fig. S1A). As plasma antibody diffusion into the brain and binding to brain GluA2 has been a hypothesized mechanism of pathology, we tested if GluA2-PLP-induced plasma IgG added to murine brain sections could bind GluA2 in the hippocampus and found plasma IgG from both Onset and Progressed mice bound natively expressed GluA2 (Fig. S1B).

**Fig. 2.**
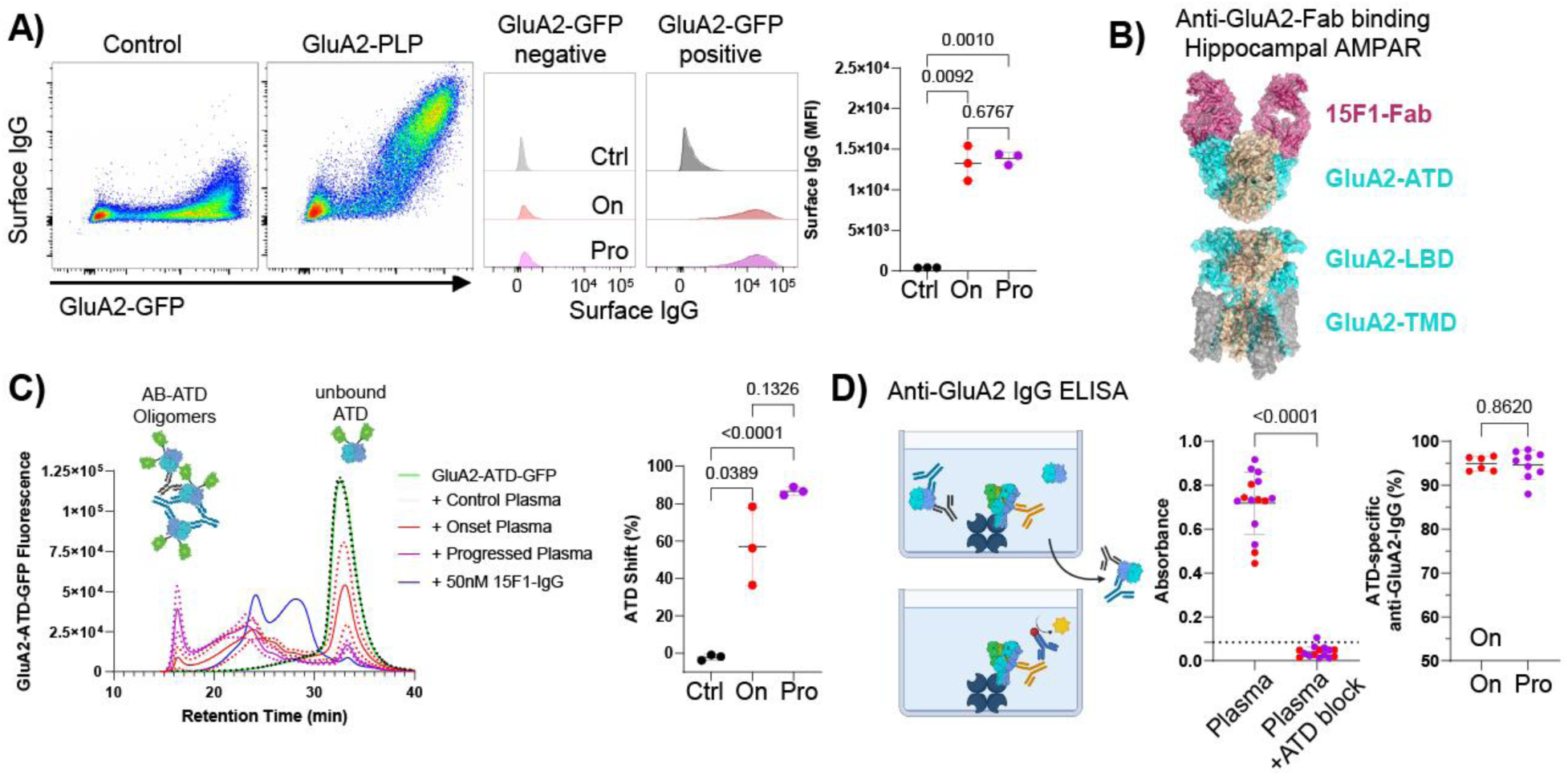
GluA2-PLP immunization elicits a robust anti-GluA2-ATD IgG response. (**A**) Plasma antibodies from Control and GluA2-PLP immunized mice binding to live cells expressing GluA2-GFP on their surface by flow cytometry (left: representative plots, right: Median fluorescence intensity (MFI) of bound IgG. (**B**) Structure of the hippocampal AMPAR bound to 15F1 Fabs (PDB ID 7LDD)^23^. (**C**) FSEC-based ATD-binding assay with mouse plasma and monoclonal anti-GluA2 IgG (clone 15F1) (left: FSEC traces, right: fraction of GluA2 ATD-GFP shifted). (**D**) Quantification of ATD-specific anti-GluA2 IgGs by GluA2-receptor ELISA ± ATD pre-incubation. Dotted line indicates the 5-sigma range (average+5*SD) of Control plasma. Schematic created with BioRender. FSEC traces are shown as mean (bold lines) ± SD (dotted line). Data presented as mean±SD. Comparison of paired samples by ratio paired t-test and groups by Welch’s t-tests. Ctrl, Control; On, Onset; Pro, Progressed.

Autoantibodies derived from AMPAR-AE patients preferentially target the ATD region^14^. Thus, we tested whether GluA2-PLP mice formed ATD-targeting autoantibodies using fluorescence-detection size exclusion chromatography (FSEC)^25^. Plasma from GluA2-PLP, but not Control mice, shifted a GluA2-ATD probe towards shorter retention times (Fig. 2C), corresponding to the larger molecular weight of probe-autoantibody complexes. Of note, plasma from GluA2-PLP mice shifted the ATD probe to shorter retention times than the control monoclonal anti-GluA2-ATD antibody, indicating the presence of polyclonal anti-ATD antibodies in GluA2-PLP plasma. To assess the proportion of ATD-specific anti-GluA2 IgG in GluA2-PLP mice, we pre-incubated plasma IgG with soluble GluA2-ATD before quantifying the antibodies still able to bind GluA2 receptor by non-denaturing ELISA. This competition resulted in an almost complete signal loss in all GluA2-PLP plasma samples in both the Onset and Progressed group indicating most GluA2-binding plasma IgG autoantibodies are ATD-specific (Fig. 2D). FSEC-competition assays with the monoclonal anti-GluA2-ATD antibody (clone 15F1) that binds the R1 lobe of the ATD revealed that some of the GluA2-PLP-induced plasma anti-GluA2-ATD IgGs also bind this site (Fig. S1C). GluA2-PLP-induced plasma IgG did not cross-react with the ATDs of GluA1 or GluA3 and only slightly with GluA4-ATD (Fig. S1D).

Next, we tested if GluA2-PLP-induced anti-GluA2 IgG reached the brain and bound to AMPARs in the hippocampus *in vivo*. Upon staining for endogenous IgGs, GluA2, and GluA1, IgG deposition was present in the hippocampus and other brain regions in most Progressed mice, but not in Onset or Control mice (Fig. 3A). The IgG deposition in Progressed mice was associated with decreased GluA2 and GluA1 immunoreactivity (Fig. 3B and 3C). High-resolution imaging in the hippocampus revealed punctate staining for GluA1 and GluA2 likely reflecting AMPAR-containing synapses in Control mice. Bright IgG puncta coinciding with reduced signal intensities of GluA1 and GluA2 puncta were detected in Progressed, but not Onset or Control mice (Fig. 3D). In Progressed mice, GluA1-puncta contained more IgG and less GluA2 and GluA1 than Onset or Control mice (Fig. 3E). This increase of IgG signal within GluA1-puncta strongly correlated with loss of GluA2-immunoreactivity, suggesting that anti-GluA2-IgGs not only reduced synaptic AMPAR expression, but also coated surface GluA2, thereby blocking staining by the monoclonal anti-GluA2-ATD antibody (Fig. 3F). In sum, GluA2-PLP immunization induced plasma anti-GluA2 IgG antibodies largely targeting the GluA2-ATD in both Onset and Progressed mice, but only Progressed mice had IgG in AMPAR-rich brain regions colocalized with GluA1- or GluA2-containing synapses.

**Fig. 3.**
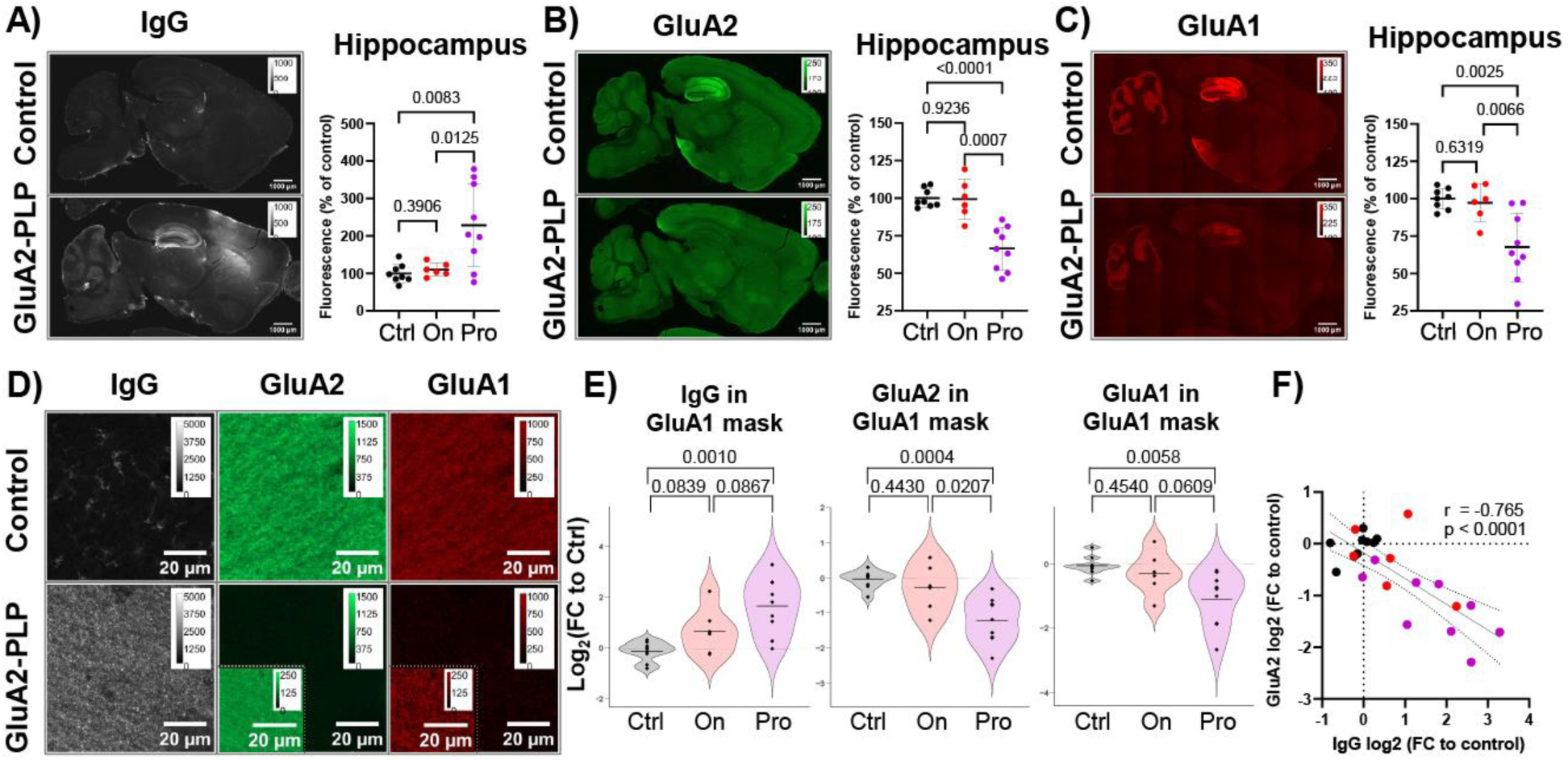
IgG deposited in the hippocampus is colocalized with AMPAR and decreased GluA1/GluA2. (**A**) Representative images of IgG (anti-mouse IgG Fc), (**B**) GluA2 (15F1 Fab) and (**C**) GluA1 (4H9 Fab) in the CNS and quantification of immunofluorescence in the hippocampus of GluA2-PLP and Control mice. Immunofluorescence was quantified as relative change to the average of Control mice (% of control). (**D**) Representative high-resolution images of punctate IgG, GluA1, and GluA2 immunoreactivity in the CA1 stratum radiatum of Control and Progressed mice. To visualize the presence of dim GluA1 puncta in Progressed mice, contrast enhanced insets (dashed line) were added. (**E**) Quantification of GluA1-colocalized IgG Fc, GluA2, and GluA1 immunoreactivity within the hippocampus. Data presented as log2-transformed fold change (FC) from the Control group. Data points represent animal means; violin contours show data distribution of synapse-level data. Groups were compared using animal means and Welch’s t-tests. (**F**) Correlation of GluA1-colocalized IgG and GluA1-colocalized GluA2-immunoreactivity colored by group (black = Control, red = Onset, purple = Progressed). Pearson correlation was performed using animal means. Data presented as mean±SD. Ctrl, Control; On, Onset; Pro, Progressed mice.

### Adaptive immune cells accumulate in the CNS with disease progression

To evaluate whether an autoimmune response against GluA2 drives neuroinflammation in mice, as seen in AE patients, we characterized the immune cells in the brains of Control and GluA2-PLP mice by flow cytometry and histology. To distinguish circulating and CNS-localized immune cells during flow cytometry, we injected mice intravenously with an anti-CD45 antibody shortly before tissue harvest to gate out circulating cells (Fig. S2A). We found increases in brain CD11b^+^CD45^mid^ microglia, CD11b^+^CD45^high^ activated myeloid cells (infiltrating monocytes and inflammatory microglia), total lymphocytes (CD45^+^CD11b^-^), CD4^+^ T cells (CD3^+^CD4^+^), CD8^+^ T cells (CD3^+^CD8^+^), non-ASC B cells (B220^+^CD138^-^ that include naïve, activated and MBCs) and ASCs (CD138^+^) in GluA2-PLP mice as compared to Controls (Fig. 4A, 4B and S2B). We use ‘ASC’ to encompass both plasmablasts (PBs) and plasma cells (PCs). We did not directly measure antibody secretion. Brain localized ASCs were almost exclusively present in GluA2-PLP mice and showed the largest fold increase.

**Fig. 4.**
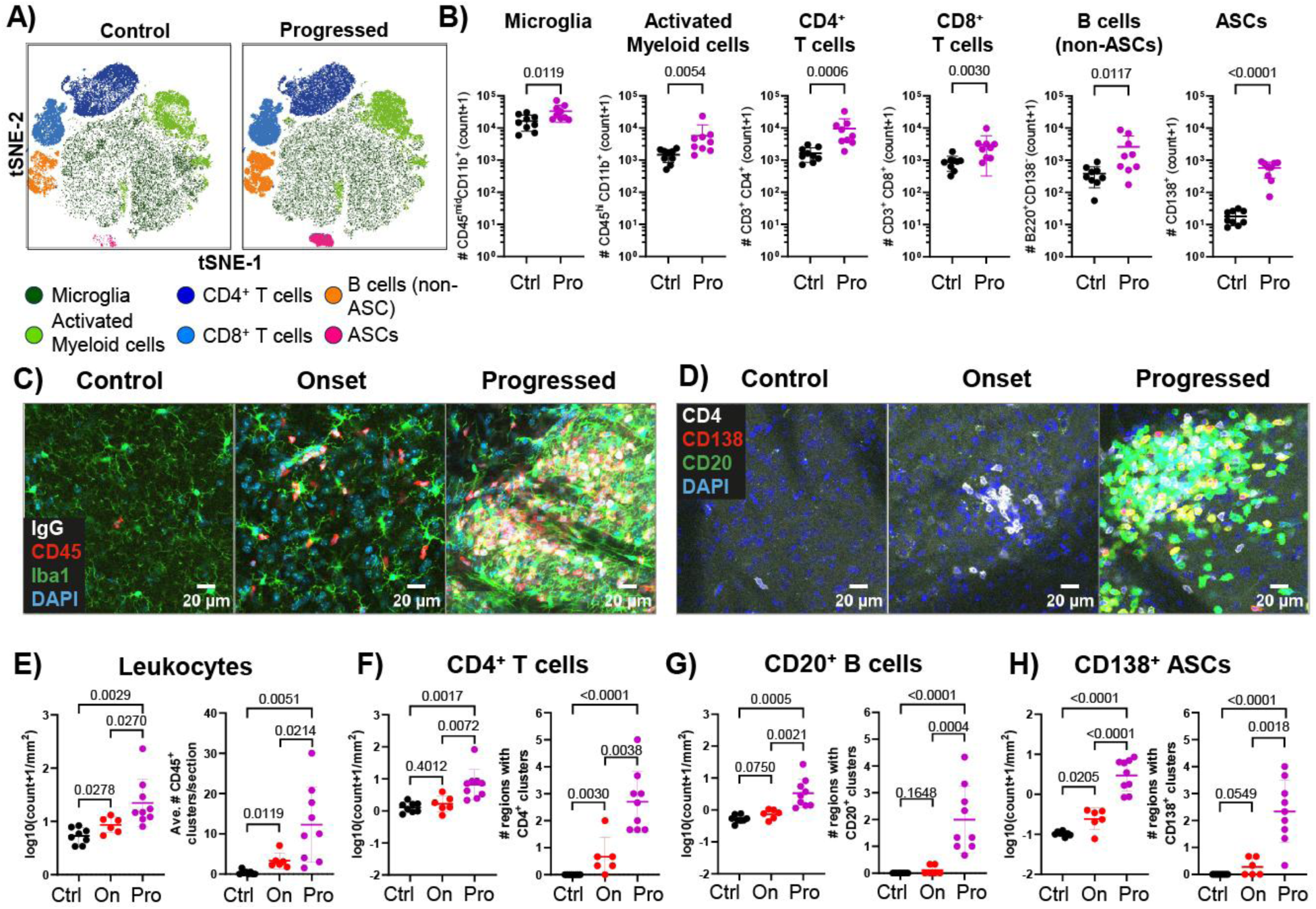
Lymphocytes accumulate in the brain during GluA2-PLP encephalitis. (**A**) Dimensionality reduction (tsne) of i.v. label^-^ immune cells pooled from brains of Control and GluA2-PLP mice and colored by cell type as defined by flow cytometry bivariate gating (n=9 mice/group, 4 experiments). (**B**) Numbers of each immune cell type in brains from Control and GluA2-PLP mice by flow cytometry (lognormal Welch’s t-test). (**C**) Representative images of microglia/myeloid cells (Iba1), antibody and ASCs (IgG), and leukocytes (CD45) in brain sections. (**D**) Representative images of immune cell infiltrates in brain parenchyma stained for CD4^+^ T cells, CD20^+^ B cells, and CD138^+^ ASCs. (**E-H**) Quantification of immune cell infiltration and clustering in brain sections of (**E**) CD45^+^ leukocytes, (**F**) CD4^+^ T cells, (**G**) CD20^+^ B cells, and (**H**) CD138^+^ ASCs (Welch’s t-tests and Mann-Whitney U tests). Data presented as mean±SD. Ctrl, Control; On, Onset; Pro, Progressed mice.

To assess the location, state, and clustering of these immune cells within the CNS, we performed histopathological examinations of whole sagittal brain sections from Control, Onset, and Progressed mice by immunofluorescence with quantification by validated Stardist-based deep learning models (Fig. S3A). All trained models showed excellent consistency with human annotations (Fig. S3B) as well as high precision, recall, accuracy, and F1 scores in image subsets of the experimental data (Fig. S3C). Brains of Control mice contained microglia with a homeostatic morphology and rare, scattered CD45^+^ lymphocytes, whereas Onset mice contained some microglia with hypertrophic soma and clusters of CD45^+^ leukocytes, both of which are characteristics of neuroinflammation. In Progressed mice, leukocyte clusters became larger, more frequent, and often contained IgG^+^ cells, likely ASCs (Fig. 4C).

Because B and T cell clustering can promote inflammation and B cell differentiation into ASCs in other tissues, we assessed CD4^+^ T cells, CD20^+^ B cells and CD138^+^ ASC accumulation and clustering in brains from Control and GluA2-PLP mice (Fig. 4D). In Control mice, lymphocyte clusters and ASCs were almost absent and only individual CD4^+^ T cells were visible. Shortly after symptom onset in GluA2-PLP mice, we observed an increased number of lymphocytes and lymphocyte clusters, that were predominantly composed of CD4^+^ T cells (Fig. 4E, 4F, and S3D). Only a few mice had mixed clusters with CD4^+^ T cells, CD20^+^ B cells and CD138^+^ ASCs (Fig. S3D). While CD20^+^ B cells were detected in similar quantities in Onset and Control in whole brain sections, CD138^+^ ASCs were enriched in Onset mice (Fig. 4G and 4H).

In the brains of Progressed mice, CD4^+^ T cells, B cells, and ASCs were increased several-fold as compared to Onset and Control mice (Fig. 4F-H). All Progressed brains contained CD4^+^ T cell-only clusters as well as numerous large mixed lymphocyte clusters in the parenchyma, in perivascular cuffs, and at the CNS borders (Fig. 4D, S3D, and S3E). These clusters appeared in more brain regions in Progressed as compared to Onset and Control mice (Fig. 4F-H), suggesting that the brain immunopathology expands with disease progression. We did not detect mature tertiary lymphoid structures^26^, frequently observed in multiple sclerosis brains^27,28^, or clusters that were primarily composed of only B cells or ASCs in any group. In sum, GluA2-PLP immunization drives CNS inflammation including innate and adaptive immune cell infiltration, lymphocyte clusters, and ASCs in the brain.

### Brain immunopathology was heterogeneous across regions, but increased in all with symptom progression

To determine if a particular set of immunopathology features or pathology in a specific brain region was consistently associated with higher clinical score for AE-like disease, we labeled the measurements of immune cells, clustering, IgG deposition, GluA2 and GluA1 immunoreactivity as pathological if they fell outside the physiological range (average +/- 3*SD) of Control mice in any of the five major AMPAR-expressing brain regions, including the hippocampus, cortex, cerebellum, striatum, and brain stem (Fig. S4A). In Onset mice, CD45^+^ and CD4^+^ clusters were common, and increases in CD45^+^ lymphocytes, CD20^+^ B cells and CD138^+^ ASCs were present in most, but not all mice (Fig. S4A). In the Progressed group, almost all parameters were pathological across all mice. A principal component analysis (PCA) of the metrically-scaled pathology parameters revealed that inter-individual and inter-regional heterogeneity increased with disease progression in two separate immunization cohorts (Fig. S4B). Because all histopathology parameters were highly correlated (Fig S4C), we combined these parameters into a pathology composite score that captured the overall immunopathology with excellent internal consistency (Cronbach’s alpha > 0.9) and positively correlated with clinical score (Fig. S4D).

To identify which brain regions were most affected by the AMPAR-AE immunopathology at onset and progression, we compared both the number of pathological findings as well as the pathology score, in each region. At symptom onset, there was heterogeneity in which brain regions were affected in each mouse (Fig. 5A), but all regions except the striatum had more pathological findings in GluA2-PLP mice compared to Control mice (Fig. 5B). However, the pathology composite score of Onset mice, which reflects the overall severity of the immunopathology, was comparable to Control mice (Fig. 5C). Upon disease progression in GluA2-PLP mice, both the number of pathological findings as well as the pathology score were increased in all brain regions compared to Onset and Control mice (Fig. 5B and 5C). The hippocampus was the most affected brain region in Progressed mice (Fig. 5B and 5C). In sum, our model of AMPAR-AE induces a spatially heterogenous brain immunopathology that spreads over multiple AMPAR-expressing brain regions with disease progression and symptom severity.

**Fig. 5.**
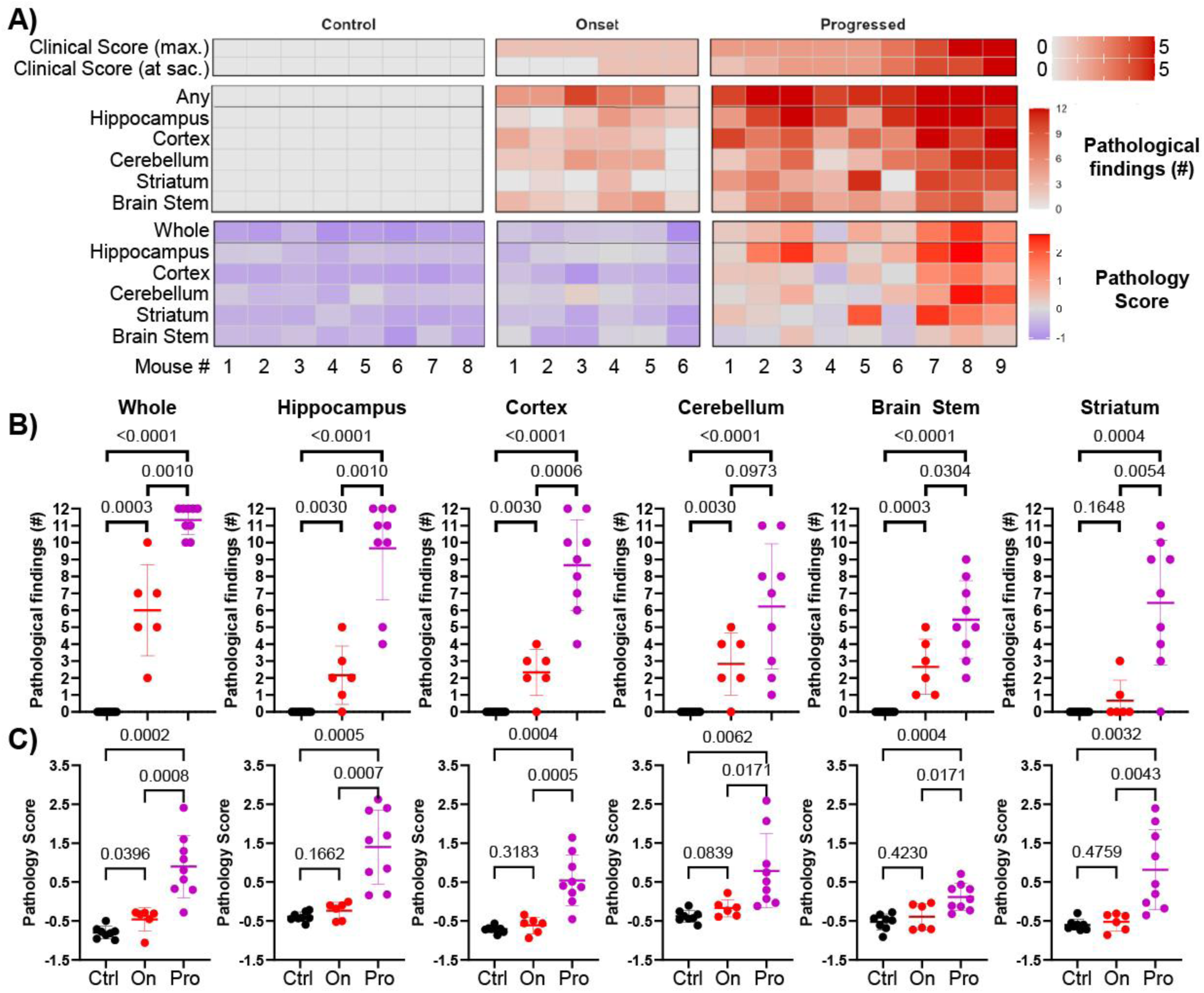
Multivariate analysis reveals spatiotemporal progression of immunopathology. (**A**) Heatmap of histopathological findings and the pathology score, a composite score that reflects the overall severity of immunopathology, in the brains of immunized mice (**B-C**) Anatomical distribution of the (**B**) number of histopathological findings and (**C**) severity of the immunopathology. Data presented as mean±SD. Welch’s t-tests and Mann-Whitney U tests. Ctrl, Control mice; On, Onset; Pro, Progressed mice.

### Plasma cells and plasmablasts in the brain parenchyma accompany symptom progression

Brain compartmentalized immune responses pose a major therapeutic challenge as the blood-brain barrier limits the CNS-access of many peripherally administered drugs^29^. Hence, we quantified ASC positioning in the parenchyma, the meninges, and the perivascular space in perfused brains of GluA2-PLP and Control mice (Fig. 6A). In Onset mice, ASCs were increased in the brain parenchyma and meninges, but not brain perivascular cuffs as compared to Control mice. ASCs further increased during disease progression with increases in all locations relative to Controls (Fig. 6A). The majority of ASCs were in the brain parenchyma, and the proportion in each location did not change with progression (Fig. 6B). Within the brain parenchyma, ASCs did not accumulate in a consistent brain region shortly after symptom onset (Fig. S5A), suggesting that localization of the initial ASCs in the brain may not depend on features of a specific brain region. In Progressed mice, ASCs increased in all brain regions but were most abundant in the hippocampus (Fig. S5B).

**Fig. 6.**
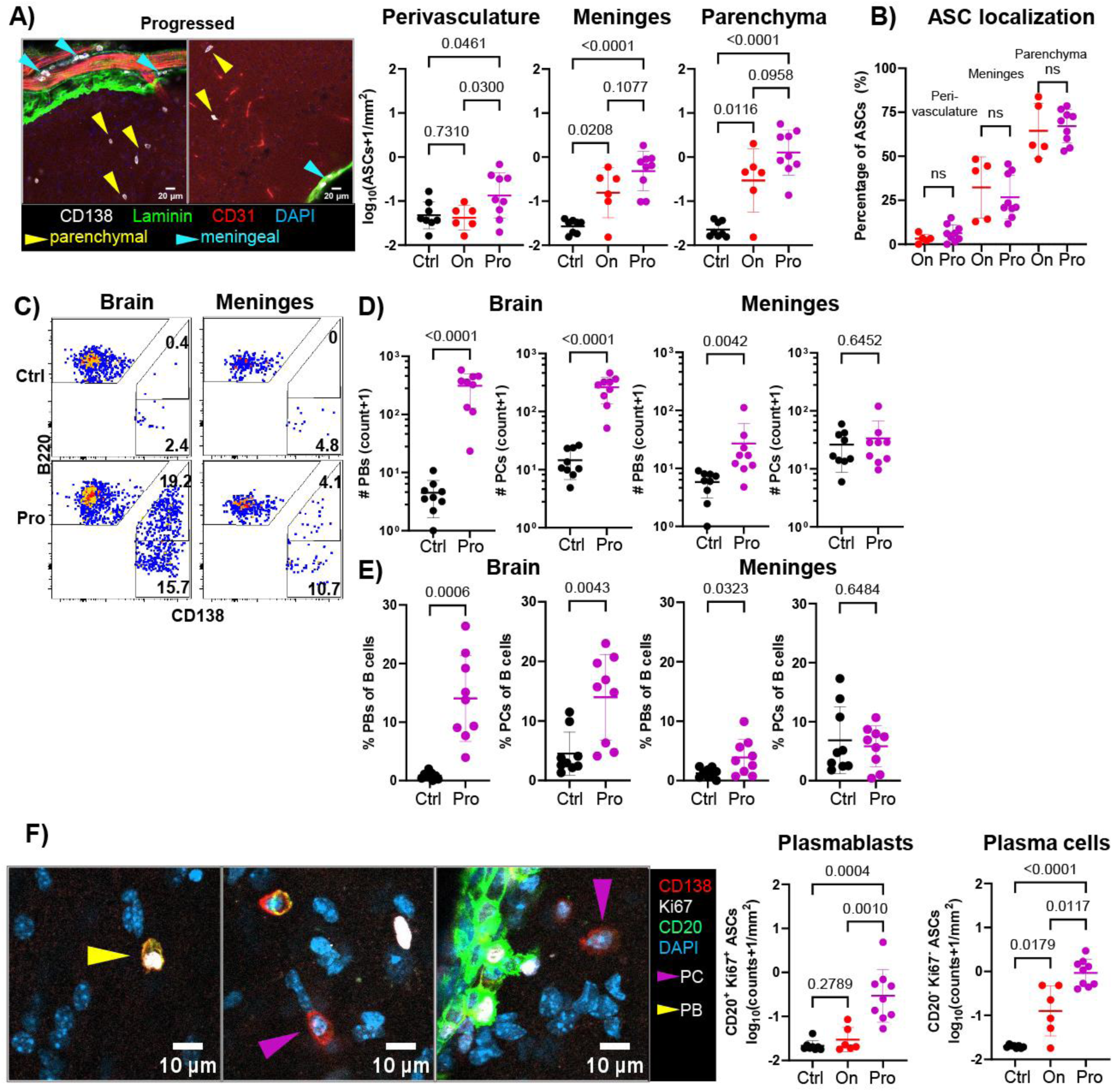
Plasma cells and plasmablasts accumulate primarily in the brain parenchyma and meninges. (**A**) Representative images and quantification of CD138^+^ ASCs across CNS compartments: perivasculature (near CD31^+^ endothelium), meninges (Laminin^+^) and parenchyma (CD31^-^Laminin^-^). (**B**) Percent of ASCs in each compartment in GluA2-PLP mice. (**C**) Representative flow cytometry gating on brain and meningeal i.v. label^-^ B cells for ASC subsets, plasmablasts (PBs, CD138^+^B220^+^) and plasma cells (PCs, CD138^+^B220^-^). (**D**) Number and (**E**) proportion of B cells that are PBs and PCs in brain and meninges of Control and Progressed mice by flow cytometry. (**F**) Representative images of plasmablasts (CD138^+^CD20^+^Ki67^+^, yellow arrow) and non-proliferating plasma cells (CD138^+^CD20^-^Ki67^-^, magenta arrow) in the brain of Progressed mice and quantification of histology. Data presented as mean±SD. Lognormal Welch’sand Welch’s t-tests. Ctrl, Control; On, Onset; Pro, Progressed mice.

ASCs are classically divided into long-lived plasma cells, that can survive and produce antibodies for months to years, and plasmablasts, which are recently generated from proliferating B cells and are often short-lived antibody producers^1,30,31^. To determine which types of ASCs were induced by GluA2-PLP immunization in the brain and meninges, we first assessed their surface and proliferation markers by flow cytometry (Fig. 6C). In contrast to brains of Control mice, that contained few PBs (CD138^+^B220^+^) and PCs (CD138^+^B220^-^), both PBs and PCs were enriched by number and proportion of the total B cell pool in the brains of Progressed mice. In the meninges, only PBs but not PCs were increased in Progressed as compared to Control mice (Fig. 6D-E). To corroborate this finding and quantify PBs and PCs in Onset brains, we developed a computational approach to phenotype ASCs in immunofluorescence images of entire sagittal brain sections. This pipeline utilized CD138^+^ ASCs masks, generated by our validated Stardist-based deep learning model, to extract CD20 and Ki67 fluorescence intensities from multi-channel immunofluorescence images, and classified ASCs into CD20^+^Ki67^+^ PBs and CD20^-^Ki67^-^ PCs by uniform bivariate gating^1,30,32^ (Fig. S6A-C). Consistent with the flow cytometry analysis, we found a pronounced increase of both PBs and PCs in the brains of Progressed versus Onset and Control mice (Fig. 6F). In contrast, only PCs, but not PBs, were increased in the brains of Onset as compared to Control mice (Fig. 6F).

### Brain ASCs synthesize AMPAR-specific autoantibodies shortly after symptom onset

Intrathecal autoantibody synthesis has been proposed as a pathomechanism in several AEs, including AMPAR-AE^1,2,33^. However, apart from a few cases in which disease-associated autoantibodies have been cloned from cerebrospinal fluid-derived B cells^34–37^, it is unknown whether autoreactive ASCs in the CNS are a source of pathogenic AMPAR-specific antibodies and where they are located. To test if brain-localized ASCs in GluA2-PLP mice are synthesizing GluA2-specific autoantibodies, we measured the ability of the antibodies inside ASCs to bind a fluorescent GluA2-ATD probe. As there were few brain ASCs in Control mice, we compared GluA2-ATD-GFP binding in GluA2-PLP immunized mice to binding in GluA1-PLP immunized mice. As with GluA2-PLP, GluA1-PLP immunization induced autoimmune encephalitis marked by hyperactivity, severe neurological symptoms, neuroinflammation, IgG deposition and loss of GluA1 immunoreactivity in the brain (Fig. S7A-D). We quantified ASCs binding the probe with our *in situ* ASC phenotyping pipeline and a GFP-only probe as GFP-binding control (Fig. S8A-B). ASCs in GluA2-PLP immunized mice specifically recognized the GluA2-ATD-GFP, but not the GluA1-ATD-GFP or GFP-only probes (Fig. S8C) as expected from the lack of cross-reactivity of GluA2 immunization-induced plasma antibodies (Fig. S1D). Furthermore, GluA2-ATD-GFP^+^ ASCs were enriched and almost exclusively detected in GluA2-immunized as compared to GluA1-immunized and liposome-immunized mice (Fig. S8C).

Using our GluA2-ATD-GFP probe, we quantified the ATD-specificity and isotype of brain-localized ASCs in GluA2-PLP mice (Fig. 7A). The proportion of brain ASCs that were ATD-specific was higher in Onset mice compared to Controls and increased further in Progressed mice (Fig. 7B). On average, the majority of ASCs in Progressed mice were ATD-specific. ATD^+^IgG^+^ ASCs, but not ATD^-^ ASCs, were enriched in Onset as compared to Control mice, indicating early presence in GluA2-PLP brains of ASCs synthesizing anti-GluA2-ATD IgG autoantibodies (Fig. 7C and S8D). The number of brain ATD^+^IgG^+^ ASCs correlated with IgG deposition in brains and clinical score (Fig. 7D). In Progressed mice, ATD^+^ IgG^-^ASCs as well as ATD^-^ ASCs were also enriched compared to Controls (Fig. S8D). Together, these findings reveal a striking immunodominance and early brain-localization of AMPAR-ATD-specific ASCs synthesizing autoantibodies which could contribute to AMPAR-AE pathogenesis.

**Fig. 7.**
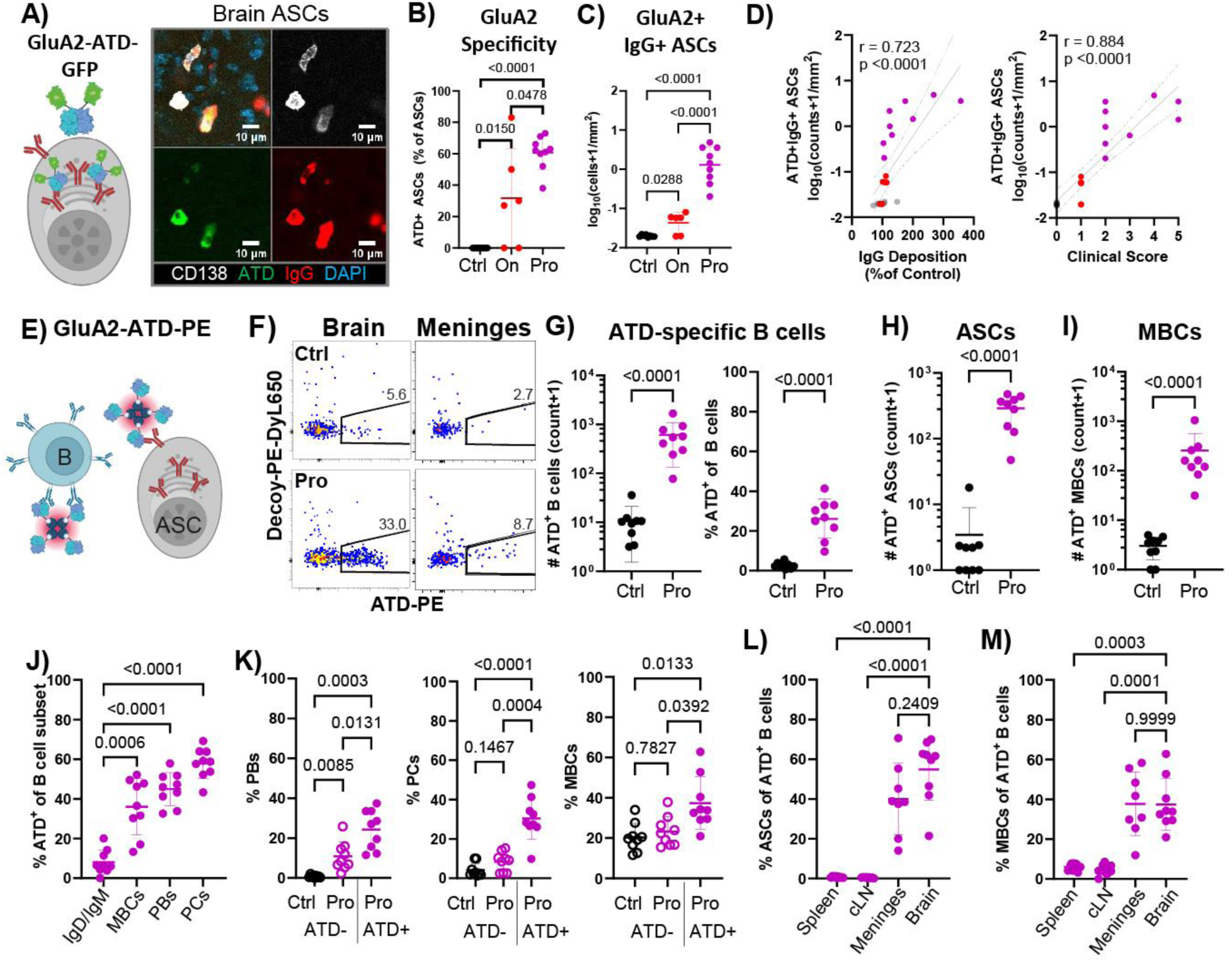
GluA2-ATD-specific ASCs and ongoing ATD-specific autoimmunity form in the brain with GluA2-PLP encephalitis. (**A-C**) Representative image, histological quantification and isotyping of GluA2-ATD-specific ASCs in brains with GluA2-ATD-GFP. (**D**) Correlation of brain GluA2^+^IgG^+^ ASCs with IgG deposition (Pearson correlation), and clinical score (Spearman correlation). (**E**) Schematic of GluA2-ATD-PE tetramer. (**F**) Representative flow cytometry gating of brain and meningeal i.v. label^-^ B cells for ATD specificity in Control and Progressed mice. (**G**) Number and proportion of ATD^+^ B cells, (**H**) number of ATD^+^ ASCs and (**I**) number of ATD^+^ MBCs in the brain. (**J**) Proportion of B cell subsets that are ATD-specific in brain. (**K**) Percentage of ATD^+^ and ATD^-^ B cells that are differentiated into PBs, PCs and MBCs in the brain. (**L-M**) Percentage of ATD^+^ B cells that are (**L**) ASCs and (**M**) MBCs across tissues of Progressed mice. Data presented as mean±SD. (B) Mann-Whitney test, (C, G-left) Welch’s t-test, (G-right, H, and I) lognormal Welch’s t-test, (J-M) Welch’s ANOVA with Dunnett’s T3 post-hoc t-test. Ctrl, Control; On, Onset; Pro, Progressed mice. Schematics created with BioRender.

### The GluA2-ATD-specific humoral autoimmune response is concentrated in the CNS during progressed GluA2-PLP encephalitis

The accumulation of autoreactive ASCs in the brain with disease severity suggests either continuous recruitment of ATD-specific ASCs formed outside the CNS and/or differentiation of ATD-specific B cells into ASCs in the brain or meninges. To track the differentiation of autoreactive ATD-specific B cells into ASCs across tissues, we built a PE-bound GluA2-ATD probe for flow cytometry (Fig. 7E). ATD-specific B cells express B cell receptors on their surface that can bind ATD. We tetramerized the GluA2-ATD to enable detection of both naïve ATD-specific B cells with low affinity B cell receptors and affinity-matured, isotype class-switched ATD-specific MBCs with high affinity B cell receptors in addition to ASCs^38^. To exclude PE-specific B cells from analysis, we also generated a decoy tetramer probe that has all of the non-ATD elements of the ATD-PE probe.

To test that our ATD-PE probe identifies ATD-specific B cells responding to intraperitoneal GluA2-PLP immunization by proliferating and differentiating into MBCs and ASCs, we analyzed probe-bound (ATD^+^) B cells in the spleens and bone marrow of GluA2-PLP and Control mice (Fig. S9A). As expected for immunogen-specific B cells, splenic ATD^+^ B cells in GluA2-PLP immunized mice were higher in number and more differentiated into MBCs than ATD^+^ B cells in Control mice or non-probe binding (ATD^-^) B cells in GluA2-PLP mice (Fig. S9B). As expected from the difference in plasma ATD-specific autoantibodies between GluA2-PLP and Control mice, bone marrow ASCs in GluA2-PLP mice had higher numbers of ATD^+^ ASCs relative to Controls (Fig. S9C). Because long-lived ASCs downregulate their B cell receptors as they persist, we tested what proportion of ATD-specific ASCs, as defined by intracellular ATD-GFP staining, still bound ATD-PE on their surface and found the majority did (Fig. S9D). Thus, the ATD-PE probe enriches autoreactive ATD-specific B cells and can be used to track them across tissues and differentiation states.

Using our ATD-PE probe, we investigated whether an ongoing ATD-specific immune response in the spleen or cervical lymph nodes, which can drain antigen from the brain, could be contributing to GluA2-PLP encephalitis. An ongoing lymphoid tissue response often includes immunogen-specific germinal center B cells differentiating into immunogen-specific ASCs and MBCs. At 10-24 days post-3^rd^ immunization, we found no difference in the number of ATD^+^ germinal center B cells or ATD^+^ ASCs in either tissue in Progressed relative to Control mice (Fig. S9E and S9F). We did find an increased number and proportion of ATD^+^ MBCs in the cervical lymph nodes as we had seen in the spleen (Fig. S9G). These ATD^+^ MBCs could contribute to the brain autoimmune response as MBCs can circulate and be recruited to inflamed tissues to rapidly proliferate and differentiate into ASCs upon encountering their antigen^26^.

To investigate if ATD-specific ASCs might be accumulating in the brain because of ongoing differentiation in the CNS, we quantified ATD-specific B cells by differentiation state in the brain and meninges (Fig. 7F and S10A). Total ATD-specific B cells were expanded by number and proportion of B cells in the brain and meninges of Progressed as compared to Control mice (Fig. 7G and S10B). This expansion included increased ATD^+^ ASCs (Fig. 7H and S10C) and increased ATD^+^ MBCs in both tissues relative to Controls (Fig. 7I and S10D). Differentiated B cells (MBCs, PBs and PCs) were more enriched for ATD-specificity than non-class switched B cells (IgD^+^ and IgM^+^) supporting antigen-specific as opposed to non-specific expansion (Fig. 7J and S10E). Indeed, the ATD^+^ B cells in the brains and meninges of Progressed mice were largely differentiated into PBs, PCs and MBCs and more so than ATD^-^ B cells in the same mice or Control mice (Fig. 7K and S10F). A slightly higher proportion of ATD^-^ B cells in Progressed mice formed PBs in the brain than controls in keeping with the histology findings and suggesting there may be a small non-ATD-specific response possibly to other epitopes on AMPAR. Finally, ATD-specific MBCs bound more ATD-PE than ATD-specific ASCs likely reflecting a difference in B cell receptor expression which could impact the efficacy of certain B cell targeted therapies (Fig. S10G). Assessing across tissues, a much higher proportion of ATD^+^ B cells in the CNS were ASCs or MBCs than in secondary lymphoid organs, suggesting ATD^+^ B cells expand and further differentiate within the CNS (Fig. 7L and 7M). In sum, pathogenic autoantibody production in the CNS is likely sustained by ongoing differentiation of GluA2-ATD-specific MBCs into PBs and PCs in the brain and meninges.

## DISCUSSION

Here we generated a mouse model of sustained and progressive anti-AMPAR autoimmune encephalitis that recapitulates key elements of the human disease to investigate the cellular underpinnings of neuropathology. Active immunization with fully assembled AMPA receptors in PLPs induced anti-AMPAR autoantibodies and progressing motor deficits. Symptom severity correlated with IgG deposition in AMPAR-rich brain regions, reduced detection of AMPAR and leukocyte accumulation in the brain parenchyma including plasma cells synthesizing AMPAR-ATD-specific antibodies. Despite the initial response in the spleen, the AMPAR-ATD-specific B cell response was amplified in the brain and meninges with progressed symptoms. Our results suggest that ongoing immune cell activity within the brain helps sustain disease activity and this model can be used to further determine the mechanisms perpetuating the cellular autoimmune response in anti-AMPAR autoimmune encephalitis. Furthermore, because of the parallels in sequence, mechanism, synaptic localization and biological function of AMPA and NMDA receptors, together with their similar autoimmune disease symptoms in humans and mouse disease models^18^, we anticipate that insights we describe in the present manuscript will also be applicable to disease mechanisms in NMDA receptor encephalitis.

Our active immunization model induced systemic anti-AMPAR autoimmunity including antibody-mediated, progressive pathology in the brain enabling study of the cellular immune response that is limited in other models. Prior rodent models of anti-AMPAR encephalitis relied on passive antibody infusion directly into the CNS, which did not induce self-sustaining pathology and could not address how and where such antibodies are generated or how they access AMPARs in the CNS^15,17^. Active immunization with glutamate receptor peptides has also been used in several AE mouse models^39–41^. For example, NMDAR-peptide immunization induced anti-GluN1-IgG but either failed to induce neuroinflammation^41–43^ or resulted in limited immune cell infiltration into the brain and only mild symptoms^39,40^. Because peptides are not conformationally-restricted, they may not duplicate surface epitopes in naturally occurring synaptic receptors in brain. In contrast, active immunization with holomeric AMPA or NMDA receptors in PLPs, causes a subacute progressive disease course with severe behavioral symptoms and immune cell infiltration in the brain parenchyma, perivascular cuffs and meninges, as observed in human NMDAR-AE cases^2,18,44–46^. These results indicate that active immunization with PLPs is an efficient tool to explore how surface receptor autoimmunity triggers autoimmune encephalitis.

Several features of PLPs may contribute to the sustained autoimmune response we observed. PLPs present intact, membrane-bound tetrameric glutamate receptors in a highly structured Y-shaped architecture, producing a repetitive display of the extracellular-facing ATD domain with conformationally-restricted epitopes. This organization likely efficiently crosslinks B cell receptors on ATD-specific B cells driving strong activation^47,48^. The repetitive and accessible display of the ATD could account for the immunodominance of ATD-specific autoantibodies in our model which is also seen in human AMPAR-AE and NMDAR-AE patients^14,18,49–52^. The fully assembled receptor also provides more CD4^+^ T cell epitopes than smaller peptides, potentially activating a larger pool of AMPAR-specific CD4^+^ T cells which can further promote the autoimmune response through inflammation and AMPAR-specific B cell activation.

Autoantibodies in the cerebrospinal fluid are postulated to perpetuate disease in AEs, including AMPAR-AE^1,2,5^, but the localization, generation and maintenance mechanisms of the ASCs producing those antibodies are largely unknown. Although in a few cases autoreactive B cells and ASCs have been identified in patient cerebrospinal fluid^33–37^, the specificity of ASCs in the meninges, or brain parenchyma of autoantibody-mediated AE cases has not been tested. Using our two AMPAR-ATD probes, we identified AMPAR-ATD-specific IgG^+^ ASCs as the only ASCs that increased in the brain parenchyma at symptom onset and further expanded in brain and meninges with progression. These included non-proliferating PCs, which could sustain autoantibody production in the CNS by surviving long-term as the inflamed CNS can produce the trophic factors required for long-lived PCs^29,32,53^.

AMPAR-specific ASCs could accumulate by recruitment, proliferation and/or local differentiation from AMPAR-specific memory B cells. While B cells and ASCs are largely excluded from the brain at homeostasis, they can accumulate during inflammation independent of antigen-specificity, either via infiltration from the meninges, the choroid plexus and cerebrospinal fluid, or via the blood-brain barrier and perivascular space^27–29,32,54,55^. At symptom onset in GluA2-PLP encephalitis mice, PCs, but not proliferating PBs, accumulated in the meninges and brain parenchyma supporting recruitment of peripherally generated PCs. At disease progression, we did not find ongoing AMPAR-specific B cell differentiation in peripheral lymphoid organs, but did in the brain and meninges. Within the CNS tissues ATD^+^ B cells were more differentiated than ATD^-^ B cells in the CNS, suggesting that they were both recruited, but only the ATD^+^ B cells encountered cognate antigen, AMPAR-ATD, and CD4^+^ T cell help, in the meninges and clusters in the brain parenchyma, that drove ATD^+^ B cell expansion and differentiation into ATD^+^ PBs. While B cell activation and differentiation in the meningeal dura-associated lymphoid tissue occurs in other neuroinflammatory diseases, whether differentiation can happen in the brain is unclear, but a possibility in our model^56^.

Coincident with elevated numbers of brain AMPAR-specific ASCs, IgG deposition colocalized with AMPARs and with reduction in GluA1/A2-immunoreactivity. GluA2-deficient Stargazer mice have a similar motor phenotype^19,20^. Thus, the behavioral alterations could be caused by autoantibody-mediated internalization of heteromeric GluA1/GluA2-receptors^23^. Furthermore, the presence of IgG puncta colocalizing with GluA1, but lack of GluA1-specific antibodies in serum, suggests that some GluA1/GluA2-containing AMPARs remain on the cell surface coated with anti-GluA2-ATD IgG autoantibodies. Coating of neuronal surface antigens by autoantibodies could facilitate neuroinflammation by initiating complement and Fc receptor signaling or direct Fc receptor-mediated internalization and degradation of AMPAR receptors by microglia contributing to behavioral phenotypes^57^.

If differentiation and survival of AMPAR-specific ASCs occurs directly within the brain parenchyma in humans as in the mouse model, this would impact the efficacy of immunosuppressive therapies to durably deplete pathogenic autoantibodies. Firstly, non-proliferating plasma cells are insufficiently targeted by current best-practice immunosuppressive therapies^1,4,30,31^. Secondly, CNS-barriers shield brain-localized immune cells from depletion by therapeutic monoclonal antibodies, even under inflammatory conditions such as in multiple sclerosis^29,58^. Thus, therapeutics that can cross into the brain and deplete memory B cells and long-lived plasma cells, such as BCMA-directed CAR T cells^59,60^, may be necessary to reduce AE recurrence and autoantibody-mediated neuropathology. Additionally, discovery of CNS-specific signals required for ASC migration and survival could provide new therapeutic targets. B cell depletion based on antigen-specificity could also improve efficacy while preserving protective humoral immunity by only depleting autoreactive B cells. The antigen probes, imaging approaches, and AMPAR-AE mouse model introduced here provide a platform to systematically evaluate such ASC-directed interventions, including promising CAAR T cell approaches^61^ that aim to eliminate B lineage cells with autoantigen-specific B cell receptors.

## Limitations of the study

First, the small sample size and heterogeneity in disease manifestation limited our analysis of pathological events at disease onset. Subtle pro-inflammatory changes that precede ASC infiltration might have been missed due to this limitation and warrant further investigation. Second, because AMPAR-AE was originally described in a predominantly female patient cohort^5^, we investigated disease mechanisms in female mice. However, larger follow-up cohort studies observed a nearly equal sex distribution in adult patients^10^. Thus, there is need for further investigation in male mice. Lastly, because we did not test current therapeutic interventions, the ability of this disease model to predict treatment responses remains to be determined. Overall, the active immunization mouse model of AMPAR-AE established here will enable future studies to determine which components of the autoimmune response contribute to disease and provides a disease-relevant framework for preclinical testing and therapeutic development.

## MATERIALS AND METHODS

### Study design

The aim of this study was to develop an active immunization model of AMPAR-AE and investigate AMPAR-specific autoimmune responses. Experiments were performed in adult female wildtype mice. AMPAR-proteoliposome- (“GluA2-PLP”) and liposome-immunized littermate (“Control”) mice were matched 1:1 for clinical scoring and flow cytometric experiments and co-housed in groups of 4 mice per cage. Allocation of immunization groups and matching was performed by random selection of co-housed mice during ear-punch identification before the start of experiments. Immunizations were performed in two independent and evenly balanced cohorts. GluA2-PLP and matched Control mice were split into two groups based on the clinical phenotype. Minimum sample size (n=6) was estimated based on similar experiments with NMDAR-PLP immunized mice^18^. Final group sizes (n=6 Onset; n=9 Progressed) were determined by the clinical disease course of GluA2-PLP mice. For the Onset group, mice were euthanized together with matched Control mice within 3±1 days after symptom onset, if the maximal clinical score did not exceed 1. The remaining mice (Progressed group and matched Controls) were monitored daily for signs of encephalitis and euthanized after reaching the humane endpoint (max. clinical score ≥ 4) or latest 8 weeks after primary immunization. As no differences were observed between Control mice matched with the Onset and Progressed groups, these Controls were pooled for data analysis. Methodological details are described in the **supplementary methods** and include statistics, use of animals, immunization, clinical scoring, organ collection, production of liposomes and recombinant proteins, as well as autoantibody assays, flow cytometry, histology, and image processing and analysis.

## List of Supplementary Materials

Supplementary Methods

Supplementary Figures S1-S10

Supplementary Data file S1

Video S1-S4

## Acknowledgments

We thank Rachel Courtney and Theresa Provitola for assistance with manuscript preparation, the OHSU Flow Cytometry Shared Resource staff for their support, the LMF City Campus at Max Planck Institute of Multidisciplinary Sciences for their support and access to computational resources, members of the Gouaux and Rodda Labs for critical discussions, Brian Jenkins of the ALM core at OHSU, and Bernard and Jennifer LaCroute for generous support. E.G. is an HHMI investigator.

## Author contributions

Conceptualization: JW, LBR, EG

Methodology: JW, GC, NYP, NS, CJS, DKS, AG, LBR

Data collection: JW, GC, NYP, LBR

Data analysis: JW, GC, LBR

Visualization: JW, GC, LBR

Funding acquisition: JW, GC, HP, LBR, EG

Project administration and supervision: LBR, EG

Writing – original draft: JW, GC, LBR, EG

Writing – review & editing: JW, GC, NS, CJS, DKS, AG, HP, GW, LBR, EG

## Data and materials availability

Data of reported analyses is provided in the manuscript, figures and the supplementary data file S1. Deep learning models as well as respective ground truth datasets, training and validation reports are available on Zenodo (https://doi.org/10.5281/zenodo.18151688).

