## Supplementary Information for "AMPAR immunization induces progressive autoimmune encephalitis with autoreactive B cells in the brain"

**Supplementary Information Table of Contents:**

Supplementary Methods

Supplementary Figures S1-S10

Supplementary Data file S1

Video S1-S4

**Supplementary Methods:**

**Statistics:** Data was analyzed and graphed using Prism (v10, Graphpad) and R (v4.4.0). Data normality was assessed using the Shapiro-Wilk test and QQ-plots. Unless stated otherwise, normally distributed data and log-normally distributed data were compared using Welch's and log-normal Welch's corrected two-sided unpaired t-tests, respectively. Ordinal or non-parametric data were compared using Mann-Whitney U tests. Spearman correlations were used to correlate data with the ordinally scaled clinical score. Unless otherwise stated, data is presented as mean ± standard deviation (SD).

**Animals:** Adult (7-8 weeks old) female C57BL/6J mice were used in reported experiments unless otherwise stated. Female BALB/c were used in initial discovery and pilot experiments. Mice were housed in standard OHSU mouse facilities, under a 12hr light/dark cycle at 75°F, 60% humidity, with ad libitum access to food and water. The OHSU Institutional Animal Care and Use Committee approved all procedures under protocol #IP00000905. Animal experiments were planned and conducted in accordance with the ARRIVE guidelines.

**Immunization:** The immunization was based on a previously established protocol for the production of monoclonal antibodies against membrane proteins[21]. Briefly, AMPAR-proteoliposomes containing 25-30µg of homotetrameric GluA2 receptors (GluA2-PLP) or homotetrameric GluA1 receptors (GluA1-PLP) per dose or equivalent amounts of empty liposomes were resuspended and adjusted to 100µL/dose with sterile saline (0.9% NaCl in water), CpG ODN 1668 (37.4µM, tlrl-1668-1, Invivogen), and MPLA (6µg/100µL, vac-mpla, Invivogen). Immunization stock solutions were freshly prepared on ice prior to each immunization. Doses were delivered close to the spleen by intraperitoneal injection into the left side of the peritoneum. Booster immunizations were performed using the same protocol at two weeks as well as one month after primary immunization.

**Clinical Scoring:** Clinical scoring was performed daily for at least 3 min per mouse as previously described for anti-NMDAR-encephalitis mice[18]. During this monitoring process the following disease stages (square brackets) were observed in AMPAR-AE mice: [1] hyperactivity characterized by excessive or disoriented response to handling in combination with hyperlocomotion and erratic movements, particularly of the head; [2] hyperactivity with episodes of behavioral quiescence, in which mouse shows body and head tremors/jitter; [3] stage 2 in combination with slight ataxia (tottering); [4] stage 2 in combination with severe ataxia (tottering + falling over) and choreiform head movements reminiscent of stargazer mice[19,20]; [4.5] stage 4 in combination with kyphosis (hunched back) while walking and resting; [5] stage 4 with lethargic episodes.

**Organ collection:** Prior to euthanasia, mice were anesthetized with isoflurane and injected retroorbitally with 2.5µg biotinylated anti-CD45 (BioLegend, clone 30-F11) to label circulating leukocytes. After 5 min, mice were sacrificed by $CO_2$ and 400 µL transcardial blood was collected into 50 µL 80mM EDTA. Lastly, mice were transcardially perfused with 20 mL 1X PBS and organs were collected. The brain was split along the longitudinal fissure into two hemispheres. One hemisphere was post-fixed for 24h at 4 °C in PBS containing 4% formaldehyde and used for histopathological experiments. The second hemisphere as well as cervical lymph nodes, spleen, and meninges were used for flow cytometry experiments. Meninges were collected from the skull using forceps. Due to the macroscopic dissection of the meninges, we do not discriminate between different meningeal layers, but this preparation usually yields a skull-attached dura mater enriched fraction, which we analyzed as 'meninges' and a brain-attached fraction enriched in leptomeninges, which we processed with the brain tissue. All tissues for flow cytometry were collected in RPMI solution (cat. #11875093) supplemented with 2% fetal bovine serum (FBS) and stored on ice until processing.

**Liposome production:** Liposomes were prepared as previously described[18,21]. Briefly, 80mg asolectin from soybean (11145-50g, Sigma), 26.7mg porcine brain polar lipid extract (141101P-500mg, Avanti), 23.3mg cholesterol (C8667-5g, Sigma), and 5mg MPLA (Lipid A, 699800P-5mg, Avanti) were dissolved in 10mL chloroform in a 100ml round bottom flask. After evaporating the chloroform, the mixture was resuspended in 10mL TBS and subjected to 15 cycles of freezing in liquid $N_2$, thawing in 37°C water bath, and 30s sonication. After the last thawing, the mixture was extruded one time through a 0.4µm filter, one time through a 0.2µm filter and 10 times through a double layer of 0.2µm filters using a 10mL Lipex thermobarrel extruder (Evonik). Afterwards, the translucent mixture was aliquoted and centrifuged for 45min at 4°C in a fixed angle rotor at approx. 150,000 g. Liposome pellets were stored at -80°C until use.

**Recombinant proteins**: The following constructs were cloned into the pEG vector[62] and expressed in tsA201 suspension cells using the BacMam expression system: (C1) rat GluA2(flip, Q607R; aa1-848 of UniprotID P19491-2) with C-terminal StrepII-tag; (C2) rat GluA2(flip, Q607R; aa1-848 of UniprotID P19491-2) with C-terminal 3C-GFP-StrepII-tag; (C3) rat GluA2-ATD (aa1-408 of UniprotID P19491) with a N406C substitution and C-terminal 3C-GFP-StrepII-tag; (C4) rat GluA1-ATD (aa1-403, UniprotID P19490) with a N401C substitution and C-terminal 3C-GFP-StrepII-tag; (C5) rat GluA1(flip; aa1-907, UniprotID P19490-2) with C-terminal 3C-GFP-StrepII-tag; (C6) human GluA1(flip; aa1-841, UniprotID P42261-2) with C-terminal 3C-GFP-StrepII-tag; (C7) human GluA2(flip, Q607R; aa1-848, UniprotID P42262-2) with C-terminal 3C-GFP-StrepII-tag. Unless otherwise stated, construct C1 (rat GluA2) was used for AMPAR-proteoliposome production and immunization. Construct C2 (rat GluA2-GFP) was used in the live cell-based assay. Construct C3 (rat GluA2-ATD-GFP) was used for *in situ* labeling of AMPAR-specific ASCs, FSEC shift assays, as well as for GluA2-ATD, and GluA2-ATD-tetramer production. GluA2-ATD was used for the ATD-block ELISA, and the GluA2-ATD-tetramer was used in flow cytometry experiments. Construct C4 (rat GluA1-ATD-GFP) and C5 (rat GluA1-GFP) were used as indicated in Control experiments. Constructs C6 (human GluA1) and C7 (human GluA2) were used for immunizations in initial discovery and pilot experiments. Expression and purification strategies were largely based on previously published methods[18,62-65] and optimized individually for each protein.

**AMPAR expression and purification:** Rat GluA2 receptors (construct C1) or rat GluA1 receptors (construct C5) were expressed in tsA201 (HEK293) suspension cells. Cells were cultured in Freestyle expression medium (12338026, Thermo) at 37 °C and 8% $CO_2$. Cells were transfected at a cell density of 2.5-3.5*$10^6$ cells/mL with baculoviruses at a multiplicity of infection (MOI) of >100. After 8-12h, 10mM sodium butyrate and 200nM ZK-200775 (#2345, Tocris) were added to the medium and the cultures were shifted to 30°C. Cells were harvested 60-72h post-infection by centrifugation and frozen at -80°C until AMPAR purification. All purification steps were performed either on ice or at 4°C. Cells were solubilized with approx. 2 volumes of solubilization buffer (20mM Tris, 150mM NaCl, 1%w/v LMNG, 0.2% CHS, 0.8 µM aprotinin, 2 µg/mL leupeptin, 2 µM pepstatin A) for 90-120min. Insoluble material was removed by ultracentrifugation and filtration through 0.45µm and 0.22µm membranes. The soluble fraction was loaded onto Strep-Tactin 4Flow® high-capacity resin (IBA-2-1250-025, Iba) packed in a gravity flow column. The resin was washed with 10-20 column volumes of purification buffer (20 mM Tris at pH 8.0, 150 mM NaCl, 0.02% LMNG) and proteins were eluted in purification buffer with 5mM desthiobiotin. The eluted protein fraction was filtered through 0.22µm filters, concentrated to 6-8mg/mL using 100kD MWCO centrifugal filter units. For the rat GluA1 construct (C5), which contained a C-terminal 3C-GFP-StrepII-tag, the GFP-tag was removed by incubating the protein at a 1:50 enzyme/protein mass ratio for 1h at 4°C with 3C protease. No tag removal was performed for the rat GluA2 construct (C1), which did not contain a GFP-tag. SEC purification was performed for both constructs in purification buffer at a flow rate of 0.5mL/min using a Superose 6 increase 10/300 gl SEC column (29091596, Cytiva). SEC-fractions containing homotetrameric AMPAR were pooled and concentrated to 1-2mg/mL using 100kD MWCO centrifugal filter units. Total protein content was estimated using absorbance at 280nm and purity was assessed by FSEC using tryptophane fluorescence and SDS-PAGE with Coomassie staining. Proper protein folding was confirmed by FSEC using conformation sensitive ATD-specific antibodies (clone 15F1 for GluA2, and clone 4H9 for GluA1). For AMPAR-proteoliposome reconstitution, freshly purified AMPAR were used. Remaining AMPAR were snap-frozen in liquid $N_2$ and stored at -80°C until use in ELISA.

**AMPAR-proteoliposome production:** Per 200µg protein, liposome aliquots corresponding to 12mg lipids were resuspended in 100-200µL FSEC buffer containing affinity- and SEC-purified homotetrameric GluA2 receptors. After adjusting the LMNG (NG310, Anatrace) concentration to 1.5mM, the mixture was nutated for 20min at 4°C. Afterwards detergent was removed by incubating the liposome-protein mix 2x for 2h and 1x overnight at 4°C with 40mg activated biobeads per 200µg protein. After biobead removal, proteoliposomes were isolated by ultracentrifugation for 45min at 4°C in a fixed angle rotor at approx. 150,000 g and stored at -80°C until use. To estimate the reconstitution efficiency, 5-10µg of AMPAR-PLPs were used for SDS-PAGE and Coomassie stained GluA2 band intensities were compared to a standard curve of purified non-reconstituted GluA2.

**ATD-GFP, ATD, and ATD-tetramer-PE production**:

Rat GluA2-ATD-N404C-3C-GFP-StrepII (short GluA2-ATD-GFP) and rat GluA1-ATD-N401C-3C-GFP-StrepII (construct C4, short GluA1-ATD-GFP) were expressed in tsA201 suspension cells transfected at a cell density of 2.5-3.5*$10^6$ cells/mL with baculoviruses encoding the construct of interest at an MOI >100. After 8-12h, 10mM sodium butyrate was added to the medium and the cultures were shifted from 37°C and 8% $CO_2$ to 30°C and 8% $CO_2$. Three days after infection, culture medium was cleared by centrifugation, neutralized with 50mM Tris base, and filtered through 0.45μm and 0.22μm filters. The medium was concentrated to around 5% of the original volume by tangential flow filtration with a 30kD filter unit. Four volumes of TBS (20mM Tris, pH 8, 150mM NaCl) were added for buffer exchange and the sample was concentrated again to around 5% of the original medium volume. Biolock (2-0205-050, Iba) was added (1mL per liter of culture) to capture residual biotin. The tangential flow concentrate was loaded onto Strep-Tactin 4Flow® high-capacity resin (1mL bed per liter of culture; IBA-2-1250-025, Iba) in gravity flow columns. The resin was washed with 10-20 column volumes of purification buffer (20 mM Tris at pH 8.0, 150 mM NaCl) and proteins were eluted in purification buffer with 5mM desthiobiotin. The eluted protein fraction was filtered through 0.22μm filters, concentrated to 2-8mg/mL using 100kD MWCO centrifugal filter units. Insoluble material was removed by ultracentrifugation and SEC was performed using a Superose 6 increase 10/300 gl SEC column (29091596, Cytiva) in purification buffer at a flow rate of 0.5mL/min. Protein fractions containing ATD-GFP were combined and concentrated using 50-100kD MWCO centrifugal filter units. Proteins were aliquoted, snap-frozen in liquid $N_2$, and stored at -80°C until further use. Tag-free ATDs for tetramer production and the ATD-block ELISA were using the 3C site between the ATD and the GFP-StrepII-tag. 3C protease (self-made in Gouaux Lab) was added to SEC-purified ATD-GFP proteins at a mass ratio of 1:50 and incubated 1h at 4°C. Tag-free ATDs were then purified in TBS by SEC using a Superose 6 increase 10/300 gl SEC column (29091596, Cytiva), concentrated to 1-2mg/mL, snap-frozen in liquid $N_2$, and stored at -80°C until further use.

For ATD-biotinylation and subsequent tetramerization with streptavidin-PE, the C-terminal cysteines were reduced by incubating 1-2mg/mL tag-free ATD in TBS with 2-10mM cysteamine (10mM for GluA2-ATD, 2mM for GluA1-ATD) for 30min at 37 °C. After reduction, cysteamine was removed by buffer exchange (2x 10-20 vol.) with 1xPBS using 30kD MWCO centrifugal filter units. Biotinylation was performed by overnight incubation of reduced ATDs (0.25-1 mg/mL) with a 20fold molar excess of EZ-Link™ Maleimide-PEG2-Biotin (A39261, Thermo) at 4°C. Free biotin was removed by buffer exchange (2x 10-20 vol.) with TBS using 30kD MWCO centrifugal filter units. Protein stability and successful biotinylation were evaluated by FSEC. For tetramer assembly, the optimal molar coupling ratios were determined by fluorescence size exclusion chromatography using 100nM biotinylated ATD combined with different molar ratios of Streptavidin-PE (#PJRS25-1, Agilent). The following ATD:streptavidin ratios were tested: 1:0, 2:1, 4:1, 8:1, 16:1, 0:1. Optimal ratios were classified as those that showed a shift of the SA-PE signal towards lower retention times (higher molecular weights) and low amounts of uncoupled ATD. Both rat GluA1-ATDs and rat GluA2-ATDs were mixed with SA-PE at a molar ratio of 4:1. After confirming the tetramer assembly by FSEC, the tetramer solution was mixed 1:1 with glycerol and stored at -20°C. Molar concentrations were estimated by dividing the moles of input SA-PE by the final volume after glycerol dilution. Decoy tetramer (CLIP-PE-Dyl650) was made as previously described from biotinylated CLIP peptide conjugated to a custom conjugated streptavidin-PE-Dylight650[66].

**Autoantibody assays:** Blood samples were centrifuged for 10 min at 5000g, and plasma fractions were stored at -20°C until use. Total anti-GluA2-receptor and anti-GluA2-ATD autoantibodies of the IgG class were quantified in mouse plasma using self-made ELISAs coated with homotetrameric GluA2 receptors. The live cell-based assay and tissue-based assay were performed using standardized IgG concentrations after purification of IgG antibodies from n=3 mice per group using protein A/G agarose (#20422, Thermo Scientific). Fluorescence-detection size exclusion chromatography (FSEC)[25] protein binding assays were performed using both crude plasma fractions as well as purified IgG.

**GluA2-autoantibody ELISA**: GluA2 autoantibodies were quantified in mouse plasma using an ELISA developed for the screening of monoclonal antibodies against natively folded membrane proteins[21]. Briefly,

Maxisorp well plates (clear, flat bottom, 96-well, #442404, Thermo) were coated overnight at 4°C with 100µL/well 3µg/mL Strep-Tactin (#2-1204-005, Iba lifesciences) in TBS. After washing with TBS, plates were blocked with 100µL of 3% bovine serum albumin (BSA) in TBS for 2h at room temperature. After blocking, plates were washed and 50µL of 2µg/mL StrepII-tagged homotetrameric GluA2 receptors in ELISA buffer (EB, 1xTBS with 0.02%w/v LMNG) supplemented with 0.2% BSA were added per well. After 1h at 4°C, plates were washed with cold EB. Plasma samples were diluted in cold EB as indicated and a 2-fold 8-step dilution series ranging from 4pM to 512pM was prepared using the 15F1 conformation-sensitive mouse monoclonal anti-GluA2 antibody[21]. Plates were incubated for 1h at 4°C with 50µL/well of diluted samples, antibody standard, or blank EB, and washed with cold EB to remove unbound antibodies. Per well, 100µL of 0.2µg/mL HRP-coupled anti-mouse IgG (#6292, ImmunoChemistryTechnologies) in EB supplemented with 0.2%BSA were added for 1h at 4°C. Lastly, plates were washed with cold EB and 100µL/well of freshly prepared detection substrate (KPL ABTS Peroxidase substrate 2-component system, 5120-0032, Sera Care) were added. Absorbance was measured after 5min at 405nm using a CLARIOstar plate reader (BMG Labtech).

To identify GluA2-autoantibody positive samples, all samples were first measured at a 1:5000 dilution. Samples were classified as positive if the absorbance values were higher than the mean+5*SD of the Control samples. For absolute quantification of GluA2 autoantibodies using a standard curve, positive plasma samples were diluted to 1:125,000. At this dilution all samples were within the linear range of the standard curve. To quantify the amount of ATD-specific GluA2 autoantibodies, positive samples were first diluted to 1:25,000, then split into two aliquots per sample which were spiked with either 2 µM GluA2-ATD or an equivalent volume of EB. After spiking, all samples were incubated for 30min at room temperature before use in the GluA2-ELISA. All samples were measured in full technical triplicates.

**Live cell-based autoantibody assay:** tsA201 suspension cells were transfected at a cell density of $2*10^6$ cells/mL with rat GluA2-GFP encoding baculoviruses at an MOI of 8. Cells were cultured at 37 °C and 8% $CO_2$ in serum-free Freestyle expression medium (12338026, Thermo). After 12h of incubation, cell culture medium was supplemented with 10mM sodium butyrate and cells were shifted to 30 °C. After 40h cells were harvested by centrifugation and resuspended at a density of $5*10^6$ cells/mL in PBS with 2% NGS for blocking. Per plasma sample, $1*10^6$ cells were aliquoted and spiked with 2µg/mL purified IgG (n=3 per group). After 30min incubation on ice, cells were washed with 10mL 1xPBS and centrifuged. Cell pellets were resuspended in 200µL 1xPBS containing 2% normal goat serum, 0.1µg/mL DAPI, and AF647-labeled anti-mouse IgG (1:1000, ab150119, Abcam). Cells were incubated once more for 30min on ice, washed with 10mL 1xPBS, centrifuged, and resuspended in 1mL 1xPBS with 2%NGS. Samples were filtered through a 70µm nylon mesh into FACS tubes. Flow cytometry data was acquired on a FACSymphony A5. After gating for live single cells, median fluorescence intensities (MFI) of the AF647 channel (surface IgG) was measured within GluA2-receptor expressing GFP positive cells and untransfected GFP-negative Control cells using FlowJo (v10).

**Tissue based autoantibody assay:** Binding of IgG, purified from plasma of AMPAR-PLP and Control mice (n=3/group), to AMPAR-rich brain regions was tested in formaldehyde-fixed 60um sagittal brain sections of a C57BL/6J wildtype mouse. Stainings were performed using standard free-floating immunofluorescence techniques. Per IgG sample, two brain sections were first permeabilized and blocked in histology buffer (HB; 1xPBS, 0.25% Triton X100) containing 2.5%NGS and then incubated for 24h at 4°C with purified 10ug/mL mouse IgG in HB and 2.5% NGS. To test if IgG autoantibodies compete with GluA2-ATD-specific monoclonal antibody Fab fragments, sections were further incubated overnight at 4°C either with HB containing 2.5%NGS or HB containing 2.5%NGS and 100nM AF647-labeled 15F1-Fab. After incubation with either 15F1-Fab or buffer only, slices were washed with HB and incubated overnight at 4°C with 1ug/mL AF555-labeled anti-Fcg (115-565-071, Jackson ImmunoResearch). Lastly, sections were washed with HB, mounted on microscopy slides, covered with AquaPolymount (18606-20, Polysciences) and 1.5H microscopy cover glasses, and imaged using the wide-field module and a 5x air objective on a Zeiss LSM980 microscope. Imaging parameters were optimized prior to the experiment and kept constant for all tissue sections. Mean fluorescence intensities of the AF647-labeled anti-GluA2-Fab and AF555-anti-Fcg-labeled IgG autoantibodies, were measured in the hippocampus using ImageJ/Fiji. IgG binding was compared across disease conditions using raw mean fluorescence intensities.

**FSEC autoantibody assays:** FSEC assays were performed on an HPLC system (Shimadzu) equipped with fluorescence detectors and a Superose 6 increase 10/300 gl SEC column (29091596, Cytiva) as previously described[21]. GFP and tryptophan fluorescence were continuously monitored for 60min per sample. For the ATD shift assay, 50nM GluA2-ATD-GFP in FSEC buffer (20mM Tris at pH 8.0, 150mM NaCl, and 0.02%w/v LMNG) were spiked with either 10µL/mL mouse plasma, 50nM 15F1-IgG (positive control), or an equivalent volume of FSEC buffer (negative control). Per sample, 70µL were injected onto the column. Samples were incubated with ATD for at least 30min prior to FSEC and kept at 4°C until injection. A constant flow rate of 0.5mL/min in FSEC buffer was employed. After FSEC, GFP signals of the unshifted GluA2-ATD-GFP peak were determined by peak integration using uniform retention time cut-offs. The ATD shift (%) was calculated by subtracting the unshifted GluA2-ATD-GFP peak area of each sample from the expected maximum signal determined from the negative Control (50nM GluA2-ATD-GFP only) and subsequent normalization to the maximum signal. Per group, ATD shifts were determined for n=3 IgG samples from n=3 mice. For the FSEC supershift assay, 10nM GluA2-ATD-GFP in FSEC buffer (20mM Tris at pH 8.0, 150mM NaCl, and 0.02%w/v LMNG) containing 3mg/mL BSA was spiked with 4µg/100µL purified mouse IgG (n=3 samples from different mice per group). In 15F1 blocking conditions, 10nM GluA2-ATD-GFP was first incubated with 40nM 15F1-Fab (anti-GluA2-ATD) before spiking with mouse IgG. Optimal 15F1-Fab blocking conditions and optimal mouse IgG concentrations, both characterized by a complete shift of free GluA2-ATD-GFP towards lower retention times, were determined prior to sample testing using dilution series ranging from 10-80nM 15F1-Fab and 1-32ug/100µL IgG from GluA2-PLP immunized mice. Void peak area was quantified by peak integration using uniform retention time cut-offs.

**Bone marrow processing for flow cytometry**: Both femurs and tibias were harvested from mice into cold RPMI with 2% fetal bovine serum (FBS). Bone marrow was released by mechanical disruption using a mortar and pestle in RPMI+2% FBS and a pipet through a 70 µm cell strainer. Cells were then centrifuged at 1500 rpm for 5 min at 4°C, the pellet resuspended for red blood cell lysis (RBC lysis, 5 mL ammonium chloride potassium (ACK) lysis buffer for 5 min) and then washed twice with 1X PBS+2% FBS (Fc buffer, FB) at 4°C. Cells were then stained with 2 pmol Decoy tetramer (CLIP–PE-DyLight650, see above), Fc

Block (BD Biosciences, anti-CD16/32, clone 2.4G2 ) and eBioscience™ Fixable Viability Dye eFluor™ 780 in FB for 10 min at room temperature followed by surface staining with 2 pmol GluA2-ATD-PE tetramer, anti-B220-BUV737 (BioLegend, clone RA3-6B2) and anti-CD138-BV650 (BioLegend, clone 281-2) for 20 min on ice. Cells were then fixed and permeabilized (Invitrogen eBioscience Transcription Factor Staining Buffer Set, cat. # 00-5523-00) for intracellular staining and stained with GluA2-ATD-GFP in permeabilization buffer for 30 min at room temperature. Cells were washed and resuspended in 1X PBS prior to flow cytometry.

**Spleen and cervical lymph nodes processing for flow cytometry**: Spleens and cervical lymph nodes were mashed through 70 µm cell strainers in RPMI+2%FBS and centrifuged for 5 min at 1500 rpm at 4 °C. Spleens were resuspended in 5 mL of ACK lysis buffer for 7 minutes at room temperature, washed, and resuspended in 5 mL RPMI+2%FBS on ice. For antigen-probe staining, cells from both tissues were pelleted and stained with 2 pmol Decoy tetramer (CLIP–PE-DyLight650, see above) in FB for 10 minutes at room temperature then moved to ice and stained with 2 pmol GluA2-ATD-PE tetramer for 20 minutes. To enrich for antigen-specific populations, the cells were washed and resuspended in 175 µL FB with 25 µL anti-PE beads (cat. #130-048-801, Miltenyi) for 30 minutes on ice. After incubation with anti-PE beads, the cells were washed in 5 mL FB and resuspended with 3 mL FB and placed on ice. The cells were run through a 70 µm filter into an LS column on the Miltenyi QuadroMACS Seperator (cat. # 130-091-051). The LS columns were washed with 1X PBS until a final volume of 12 mL was reached. Subsequently, the LS column was removed from the separator and placed on a 15 mL conical tube and the fraction was eluted using 5 mL of 1X FB. This fraction was then washed and resuspended with Fc Block (BD Biosciences, anti-CD16/32, clone 2.4G2 ) and eBioscience™ Fixable Viability Dye eFluor™ 780 in FB for 10 min at room temperature. The following antibodies were used to phenotype lymphocytes from secondary lymphoid organs: anti-CD45-biotin (BioLegend, clone 30-F11), anti-streptavidin-BV786 (BD Biosciences, Cat. No. 563858 ), anti-IgD-BUV395 (BD Biosciences, clone 11-26c.2a), anti-B220-BUV737 (BioLegend, clone RA3-6B2), anti-CD3-BV510 (BioLegend, clone 17A2), anti-CD138-BV650 (BioLegend, clone 281-2), anti-IgM-BV711 (BioLegend, clone RMM-1), anti-GL7-AF488 (BioLegend, clone GL7), anti-CD38-

AF700 (BioLegend, clone 90), eBioscience™ Fixable Viability Dye eFluor™ 780. Following staining, cells were fixed in 1% PFA then washed and resuspended in 1x PBS prior to flow cytometry.

**Brain and meninges processing for flow cytometry**: Half of the brain was collected for flow cytometry in RPMI+2%FBS on ice and subsequently minced and mashed through a 70 µm cell strainer. 3 mL of SIP percoll (2.7 mL percoll + 0.3 mL 10X PBS) was added to each sample and samples were centrifuged for 30 minutes at 1000g at room temperature. The myelin layer and supernatant were removed, and the cell pellet was resuspended in 1x PBS and filtered through a 70 µm cell strainer before centrifuging for 10 min at 4°C at 600g and resuspending in FB. Meninges were mashed through a 70um strainer and centrifuged 5 min at 1500 rpm at 4 °C before resuspending for antibody staining. Cells were stained with 2 pmol Decoy tetramer (CLIP–PE-DyLight650, see above), Fc Block (BD Biosciences, anti-CD16/32, clone 2.4G2 ) and eBioscience™ Fixable Viability Dye eFluor™ 780 in FB for 10 min at room temperature followed by surface staining with 2 pmol GluA2-ATD-PE tetramer (see above), anti-IgD-BUV395 (BD Biosciences, clone 11-26c.2a), anti-CD4-BUV486 (BD Biosciences, clone GK1.5), anti-B220-BUV737 (BioLegend, clone RA3-6B2), anti-CD45.2-BV421 (BioLegend, clone 104), anti-CD3-BV510 (BioLegend, clone 17A2), anti-CD8a-BV605 (BioLegend, clone 53-6.7), anti-CD138-BV650 (BioLegend, clone 281-2), anti-IgM-BV711 (BioLegend, clone RMM-1), anti-CD11b-AF488 (BioLegend, M1/70) and anti-streptavidin-BV786 (BD Biosciences, Cat. No. 563858) for 20 min on ice. Following staining, cells were fixed in 1% PFA, washed, and resuspended in 1X PBS prior to flow cytometry.

**Flow cytometry and data analysis:** Prior to the collection of each sample, 5 µL of CountBright Absolute Counting Beads (cat. #C36950) were added to determine absolute cell counts. Brains, meninges, spleen, cervical lymph nodes and bone marrow cells were run on the BD FACS Symphony A5. Data were analyzed using FlowJo v.10 (Becton Dickinson) software. Flow cytometry data were analyzed using FlowJo v.10. For dimensionality reduction, data from n = 9 mice per group (Control and Progressed) were first gated to include live CD45 intravenous label negative cells. Events from all mice within each group were then concatenated into a single dataset per group to enable group level visualization while minimizing mouse-to-mouse variability. The t-distributed stochastic neighbor embedding (t-SNE) was performed in FlowJo v.10 using the following parameters as input dimensions: CD4, IgD, B220, CD45.2, CD3, CD8a, CD138, IgM, and CD11b. The default FlowJo tSNE settings were used unless otherwise specified. Clusters identified in tSNE dimensions were subsequently annotated and defined based on our established bivariate gating strategies applied to the original parameters, ensuring consistency with conventional manual gating approaches. Flow cytometry experiments were performed across five independent experimental days for both CNS and secondary lymphoid organ tissues. Absolute cell numbers were determined using counting beads. Cell counts followed a log-normal distribution and are therefore presented on a log scale of (counts + 1). Brain cell numbers represent cells isolated from one-half of a brain (other half used for histological analysis) and were not extrapolated to whole brain values. For percentage-based quantification, parent populations were required to contain a minimum of 10 events for inclusion.

**Tissue processing for histopathology**: Immunofluorescence staining of formaldehyde-fixed tissues were performed in 60um sagittal brain sections using standard free-floating IHC techniques. Unless otherwise stated, tissue sections were permeabilized with 0.5% Triton X-100 in PBS and blocked with 5% normal goat serum (NGS) for 1h at room temperature. Afterwards, tissue sections were incubated either in parallel or sequentially with fluorescently labeled antibodies. During the last antibody incubation step, DAPI was added at a concentration of 0.1µg/mL to counterstain nuclei. After each incubation, tissue sections were washed at least 3x15min in 2mL HB. Sections were mounted on microscopy slides and covered with AquaPolymount (18606-20, Polysciences) and 1.5H microscopy cover glasses. The following antibody panels were used:

P1, AMPAR and autoantibody panel: primary staining for ~48h with AF647-labeled anti-GluA1-Fab (100nM, clone 4H9, produced in the Gouaux lab[21,67]), GFP-tagged anti-GluA2-Fab (100nM, clone 15F1, produced in the Gouaux lab[21,22]), AF555-labeled goat anti-Fcg (100nM, 115-565-071, JIR).

P2, basic neuroinflammation panel: primary staining for ~48h with rabbit anti-Iba1 (1:400, 019-19741, Wako) and rat anti-CD45 (1:400, clone 30F11, 103102, Biolegend); secondary staining for ~36h with AF488 goat anti-rabbit IgG (1:1000, ab150081, Abcam), AF568 anti-rat IgG (1:1000, A11077, Thermo), AF647 anti-mouse IgG (1:1000, ab150119, Abcam).

P3, lymphocyte panel: primary staining overnight with rabbit anti-CD20 (1:400, clone E3N70, 701685, Cell Signaling); secondary staining overnight with AF594 anti-rabbit IgG (1:1000, ab150080, Abcam), AF647 anti-CD138 (1:400, clone 281-2, 142526, Biolegend), AF488 anti-CD4 (1:400, clone RM4-5, 100529, Biolegend).

P4, ASC antigen-specificity panel: primary staining overnight with AF594 anti-mouse IgG (1:400, cross-adsorbed to rat IgG, ab150120, Abcam), and GluA2-ATD-GFP (10μg/mL, produced in the Gouaux lab); secondary staining overnight with AF647 anti-CD138 (1:500, clone 281-2, 142526, Biolegend), and GluA2-ATD-GFP (10μg/mL, produced in the Gouaux lab).

P5, ASC proliferation status panel: primary staining overnight with rabbit anti-CD20 (1:400, clone E3N70, #701685, Cell Signaling), and rat anti-Ki67 (1:400, clone SolA15, 14-5698-82, eBioscience); secondary staining overnight with AF488 anti-rabbit IgG (1:1000, A11008, Invitrogen), and AF568 anti-rat IgG (1:1000, A11077, Invitrogen); tertiary staining overnight with AF647 anti-CD138 (1:400, clone 281-2, 142526, Biolegend).

P6, ASC localization panel: primary staining overnight with rabbit anti-laminin (1:1000, ab11575, Abcam); secondary staining overnight with AF488 anti-rabbit IgG (1:1000, A32731, Invitrogen), AF594 anti-CD31 (1:200, clone MEC13.3, 102520, Biolegend), and AF647 anti-CD138 (1:400, clone 281-2, 142526, Biolegend).

P7, AMPAR-specific B cell localization panel: primary staining overnight with rabbit anti-CD20 (1:400, clone E3N70, 701685, Cell Signaling); secondary staining overnight with AF594 anti-CD31 (1:200, clone MEC13.3, 102520, Biolegend), AF647-labeled anti-rabbit IgG (1:1000, ab150083, Abcam), GluA2-ATD-GFP (10μg/mL, produced in the Gouaux lab).

P8, T cell localization panel: primary staining overnight with rabbit anti-laminin (1:1000, ab11575, Abcam); secondary staining overnight with AF647 anti-rabbit IgG (1:1000, ab150083, Abcam), AF594 anti-CD31 (1:200, clone MEC13.3, 102520, Biolegend), and AF488 anti-CD4 (1:400, clone RM4-5, #100529, Biolegend).

P9, T cell interaction panel: primary staining for ~36h with rabbit anti-Iba1 (1:500, 019-19741, Wako); secondary staining overnight with AF594 anti-rabbit (1:1000, ab150080, Abcam), AF488 anti-CD4 (1:400, clone RM4-5, #100529, Biolegend), and AF647 anti-MHCII (1:400, anti I-A/I-E, clone M5/114.15.2, 107618, Biolegend).

**Image acquisition and data processing for histopathology:** Image acquisition was performed on a LSM980 microscope (Zeiss). Acquisition parameters were optimized for each experiment and kept constant within each experiment. For the AMPAR and autoantibody panel P1, whole sagittal brain slices were first imaged using a wide-field module, a 5x air objective (NA 0.16) and the following excitation and detection wavelength: GFP-channel excitation 450-490nm, detection 500-555nm; AF555/594 channel excitation 538-562nm, detection 570-640nm; AF647 channel excitation 625-655nm, detection 665-715nm. Additional high-resolution images of individual synapses within hippocampal subregions were acquired using the Airyscan 2 module (Zeiss), a 63x oil objective (NA 1.4), a pixel scaling of 42.5nm, and the following excitation and detection wavelengths: GFP-channel excitation 488nm, detection 495-548nm; AF555/594 channel excitation 561nm, detection 574-627nm; AF647 channel excitation 631nm, detection 659-735nm. Per animal, three images were acquired within the CA1 stratum radiatum, three images were acquired within the dentate gyrus molecular layer, and two images were acquired within the CA2/CA3 stratum radiatum. All other antibody panels (P2-P9) were imaged using the Airyscan 2 module, a 20x air objective (NA 0.8), a pixel scaling of 460nm and the following excitation and detection wavelengths: DAPI channel excitation 405nm, detection 422-477nm; GFP/AF488 channel excitation 488nm, detection 499-548nm; AF555/561/594 channel excitation 561nm, detection 574-627, AF647 channel excitation 631nm, detection 659-735nm. Whole sagittal images were acquired as tile scans and stitched using the DAPI channel as a reference. Airyscan image processing was performed in Zen Blue (Zeiss) using default parameters. Regions of interest included the hippocampus, cortex, striatum, cerebellum, and brain stem and were segmented according to the Allen Mouse Brain reference atlas using QuPath v0.5.0[68] and Fiji/ImageJ version 2.14.0/1.54f[69]. For quantification of IgG autoantibody deposition, GluA1-, and GluA2-immunoreactivity, mean fluorescence intensities were determined within each brain region of interest in widefield microscopy images using ImageJ. To account for brain region specific differences in AMPAR expression, the data was normalized to the respective region average of Control mice. Clusters of lymphocyte populations (≥ 10 cells in close proximity), were manually quantified per region of interest.

For quantification of cells and synapses, Stardist[70] based deep learning models were used to segment puncta in GluA1 single channel images, lymphocytes in CD45, Iba1, and DAPI triple channel images, CD4+ T cells in CD4 single channel images, CD20+ B cells in CD20 single channel images, and CD138+ ASCs in CD138 single channel images. Models were trained and validated on independent ground truth datasets generated by manual annotation of a randomly selected fraction of the experimental data as well as pilot data. Training and validation were performed using modified ZeroCostDL4Mic jupyter notebooks[71]. Ground truth datasets, training parameters, deep learning models, as well as the training and validation reports are available on Zenodo (https://doi.org/10.5281/zenodo.18151688). Cell counts were normalized to the area of the quantified region of interest. As cell counts and densities of lymphocyte populations ranged over several orders of magnitude, they were log-transformed for group comparisons as indicated in the graphs. Per animal and cell type, three sagittal brain sections were analyzed, and data was averaged within each mouse and brain region. CD45 positive lymphocytes were quantified using panel P2 and three technical replicates per mouse. CD20 positive B cells were quantified using CD20 single channel images of panel P3, P5, and P7. CD138 positive ASCs were quantified using CD138 single channel images of panel P3, P4, and P6. CD4 positive T cells were quantified in CD4 single channel images of panel P3, P8, and P9.

To construct a composite score that summarizes the histopathological findings, the log-transformed cell densities ((count+1)/mm$^2$) of CD45$^+$ lymphocytes, CD20$^+$ B cells, CD4$^+$ T cells, and CD138+ ASCs, as well as the control-normalized IgG deposition, GluA1-, and GluA2-immunoreactivites were z-transformed and averaged after adjusting for directionality of the effect. The internal consistency of the composite score was evaluated using Cronbach’s alpha. Furthermore, Z-transformed values were used for principal component analysis (PCA) to visualize and summarize the multivariate structure of the histopathological parameters across brain regions and disease conditions. For animal-wise PCA, data from different brain regions was averaged within each animal after Z-transformation.

To score histopathological events (1=present, 0 = absent) individually within each animal and each brain region, reference values (mean, standard deviation, and 3σ range) were obtained for each region of interest from liposome immunized Control mice. Lymphocyte clusters, lymphocyte counts, and IgG deposition were classified as pathological if they were above the 3σ range (≥mean+3*SD) of Control mice. GluA1 (clone 4H9) and GluA2 (clone 15F1) immunoreactivity were classified as pathological if they were below the 3σ range (≤mean-3*SD) of Control mice. Per animal, histopathological parameters were classified as positive if at least one ("Any") brain region was pathological.

To assess IgG, GluA1, and GluA2 fluorescence intensities within GluA1-containing synapses, GluA1-puncta segmentations were overlayed with the original images to extract the mean fluorescence intensities of respective fluorescence channels for each individual synapse using ImageJ. For quality control measures, i.e. to control for random colocalization due to crowding, background fluorescence was sampled by rotating the image 180° relative to the cell/puncta segmentations ("flip control"). To evaluate synaptic IgG, GluA2, and GluA1 signals across disease conditions, log-transformed mean fluorescence intensities of individual synapses were averaged within each animal. Groups were compared using animal averages and Welch's corrected t-tests.

To determine cellular phenotypes of individual ASCs, ASC segmentations were used to extract mean fluorescence intensities of the Ki67 and CD20 channels in panel P5 as well as the IgG and ATD-GFP channels in panel P4. Data was extracted for each ASC as described for GluA1-puncta using correctly oriented images and 180° rotated images. Flip controls were used to determine positivity thresholds for each marker. Fluorescence positivity thresholds were kept constant across all animals and, analogous to flow cytometry, used to classify ASC phenotypes by bivariate gating.

**Supplementary Figures and Legends:**

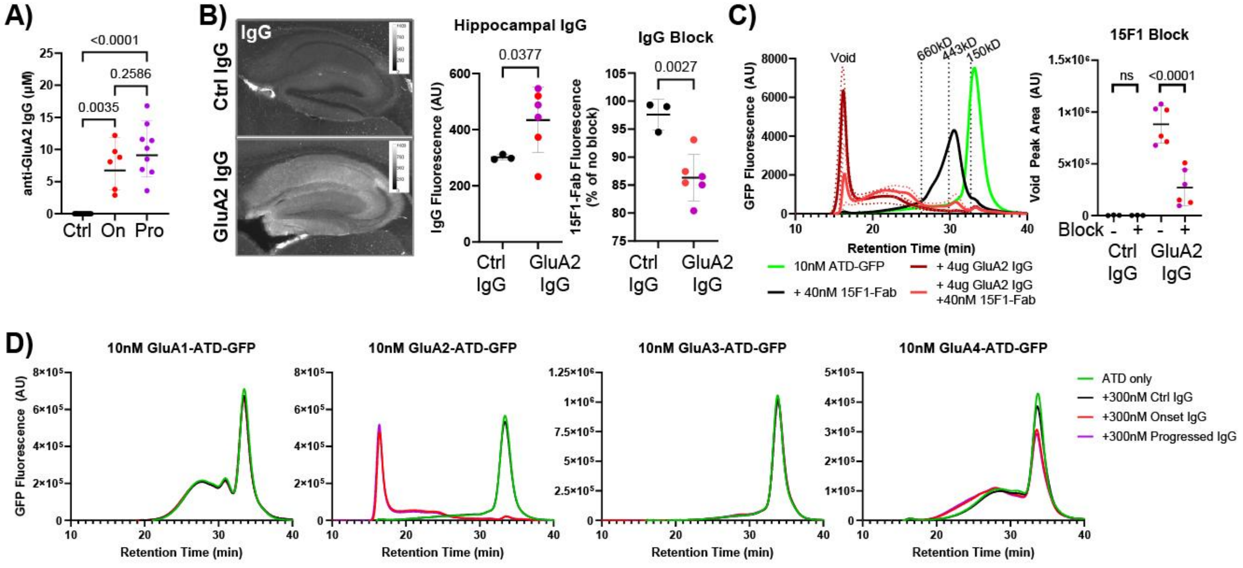


**Fig. S1. Characterization of GluA2-autoantibody binding.** (**A**) Quantification of plasma anti-GluA2 IgG autoantibodies by non-denaturing ELISA (Ctrl, Control; On, Onset; Pro, Progressed). (**B-C**) Competition of Control (Ctrl) and GluA2-PLP (GluA2) plasma IgG with anti-GluA2 antibody (clone 15F1) assessed using (**B**) a tissue-based assay with IgG block and (**C**) an FSEC supershift ATD binding assay with 15F1-Fab block. (**D**) Cross-reactivity of plasma IgG to GluA1-GluA4 ATDs determined by FSEC. FSEC traces are shown as mean (bold lines) ± SD (dotted line). Data presented as mean±SD. Groups were compared using Welch's t-tests and paired samples (C) were compared using a ratio paired t-tests.

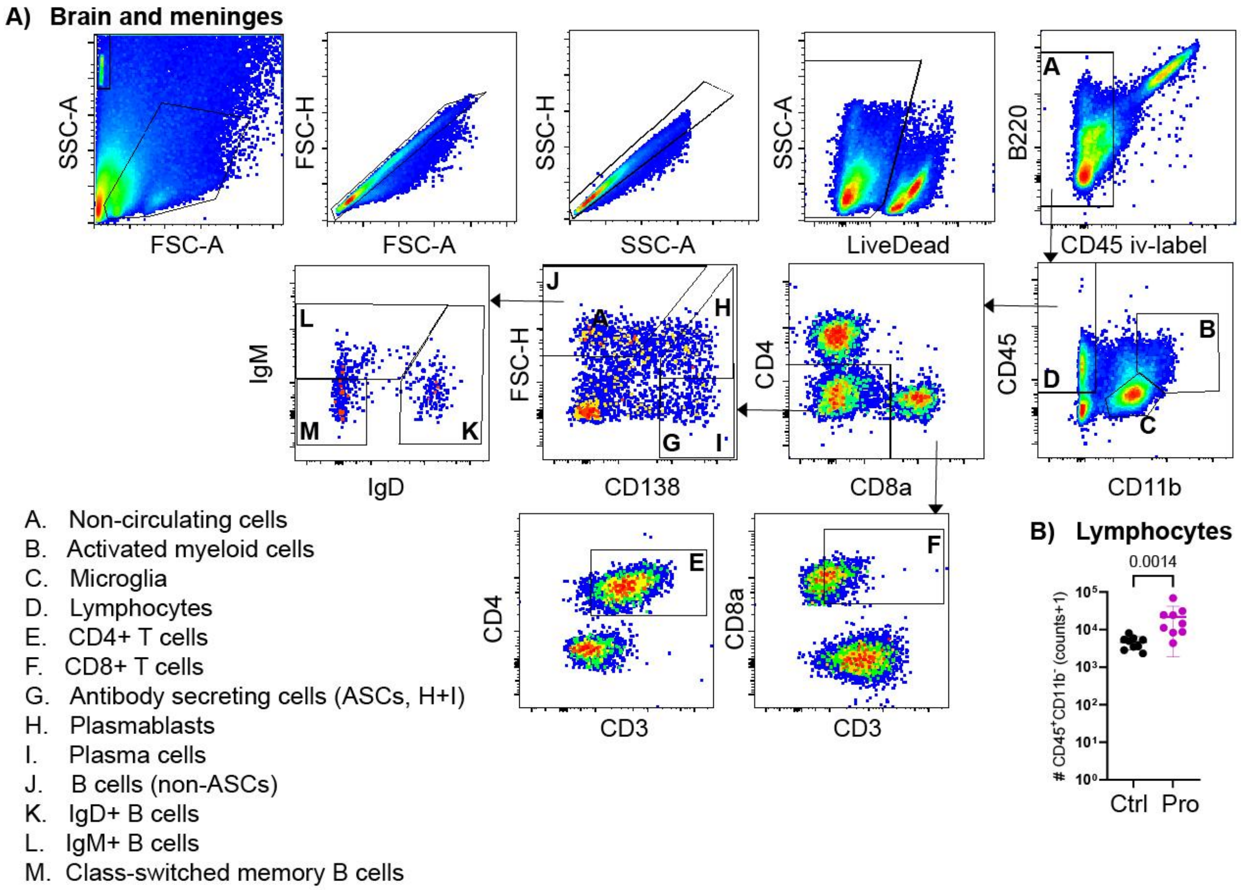


**Fig. S2. Flow cytometric quantification of immune cells in the CNS.** (**A**) Flow cytometry gating scheme used for brain and meninges (brain shown) with immune subsets indicated. (**B**) Numbers of lymphocytes (i.v. label$^-$CD45$^+$CD11b$^-$) in brains from Control and GluA2-PLP Progressed mice (mean ± SD, lognormal Welch's t-test, 4 experiments). Ctrl, Control; Pro, Progressed.

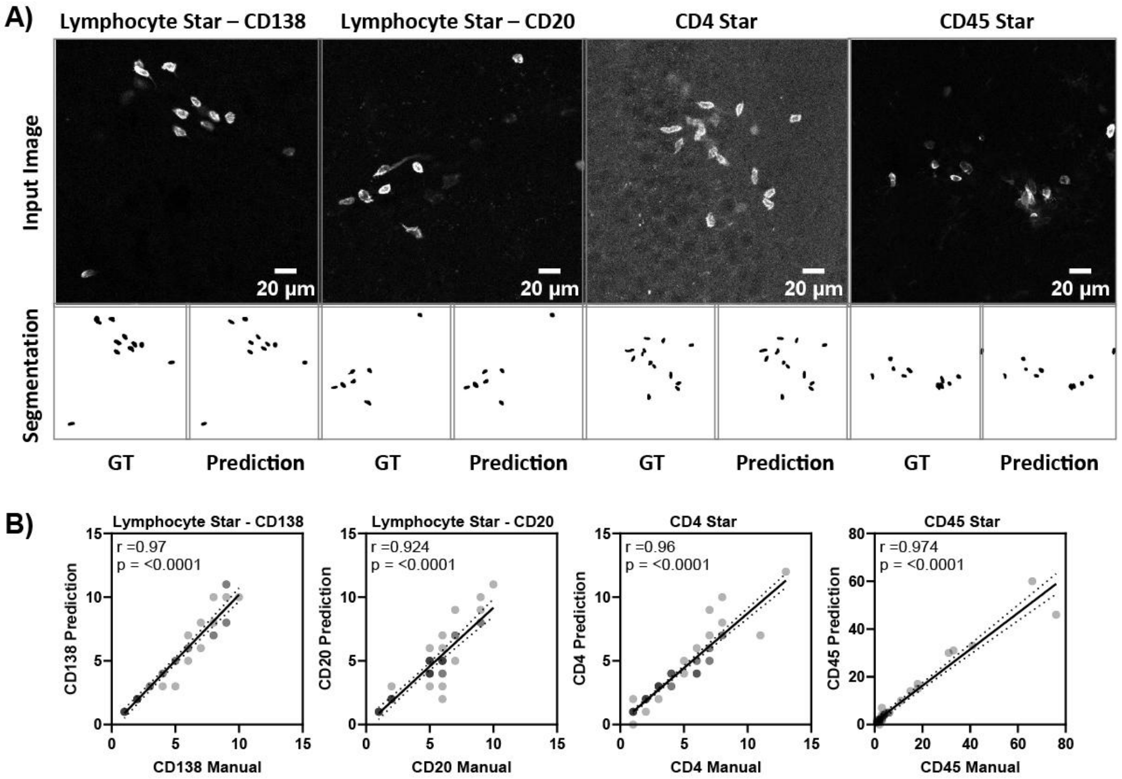


C)

| Model | Lymphocyte Star | | CD4 Star | CD45 Star |
|---|---|---|---|---|
| Image channels | CD138 | CD20 | CD4 | CD45, Iba1, DAPI |
| N (images; cells) | 42; 182 | 50; 217 | 94; 259 | 34; 377 |
| Reliability/Correlation Human vs Model (r) | 0.971 | 0.924 | 0.960 | 0.975 |
| Precision | 0.97±0.06 | 0.94±0.11 | 0.97±0.12 | 0.89±0.22 |
| Recall | 0.96±0.09 | 0.88±0.16 | 0.93±0.15 | 0.86±.22 |
| Accuracy | 0.93±0.10 | 0.84±0.19 | 0.92±0.17 | 0.81±0.24 |
| F1 score | 0.96±0.06 | 0.90±0.13 | 0.95±0.13 | 0.87±0.21 |

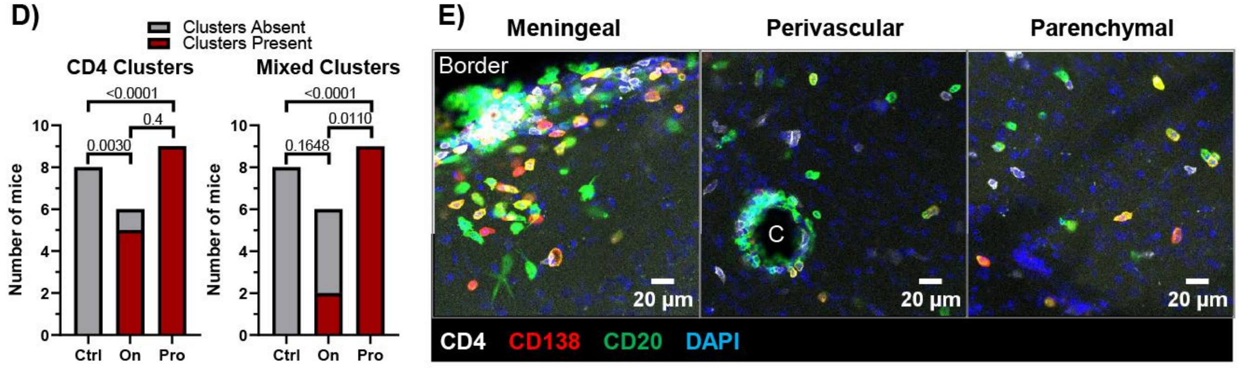


**Fig. S3. Histological quantification of immune cells in the brain. (A)** Example images of the input images, the ground truth (GT) segmentation, and the prediction. (**B**) Inter-rater-reliability determined by Pearson correlations using human annotations vs DL-model predictions. (**C**) Validation parameters presented as mean ± SD of images in the validation dataset. GT, ground truth. (**D**) Numbers of mice with (red) and without (gray) CD4+ T cell clusters and mixed lymphocyte clusters in the brains of Control, Onset and Progressed mice (Fisher's exact test). (**E**) Representative images of immune cell infiltrate in brain sections of Progressed mice stained for $CD4^+$ T cells, $CD20^+$ B cells, and $CD138^+$ ASCs. Border, cortical brain border covered by leptomeninges; C, perivascular cuff. Ctrl, Control; On, Onset; Pro, Progressed.

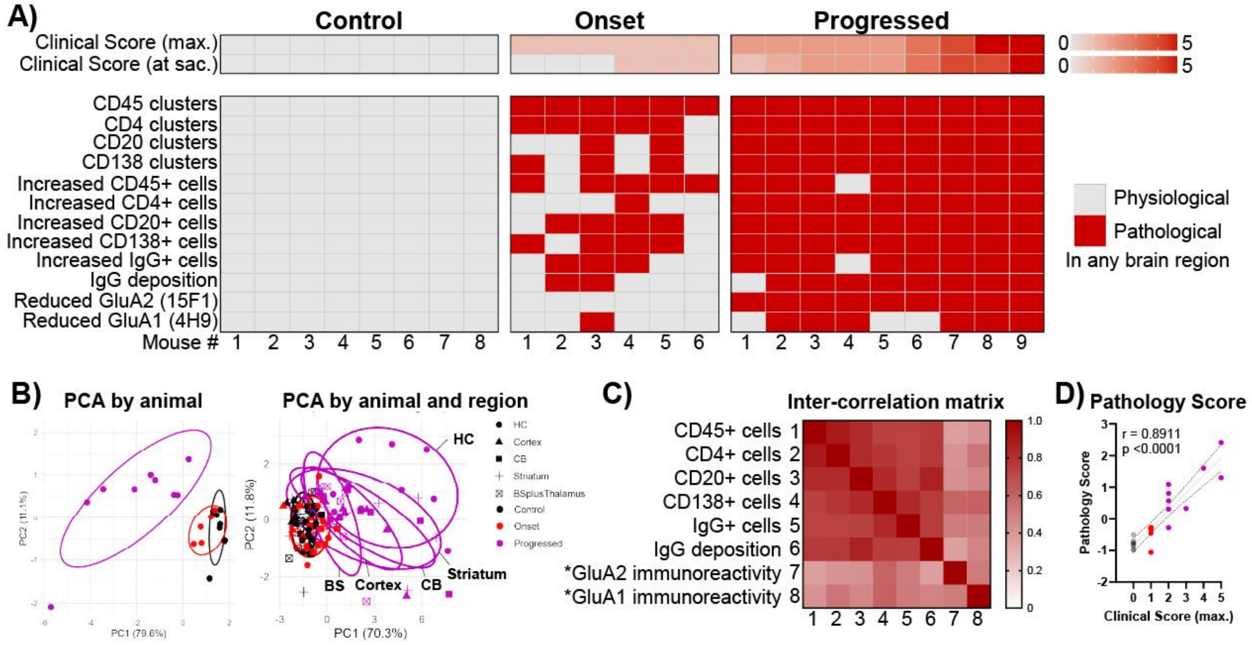


**Fig. S4. Histopathological assessment and construction of pathology composite score.** (**A**) Heatmap of histopathological findings in the brains of immunized mice. Parameters were classified as pathological (red) if any brain region had findings outside of the physiological range defined by the average ± 3*SD of Control mice. (**B**) Principal component analysis (PCA) of metrically-scaled histopathological parameters. (**C**) Inter-correlation matrix of histopathological parameters. *, directionality-aligned parameters. (**D**) Spearman correlation of clinical score with pathology composite score.

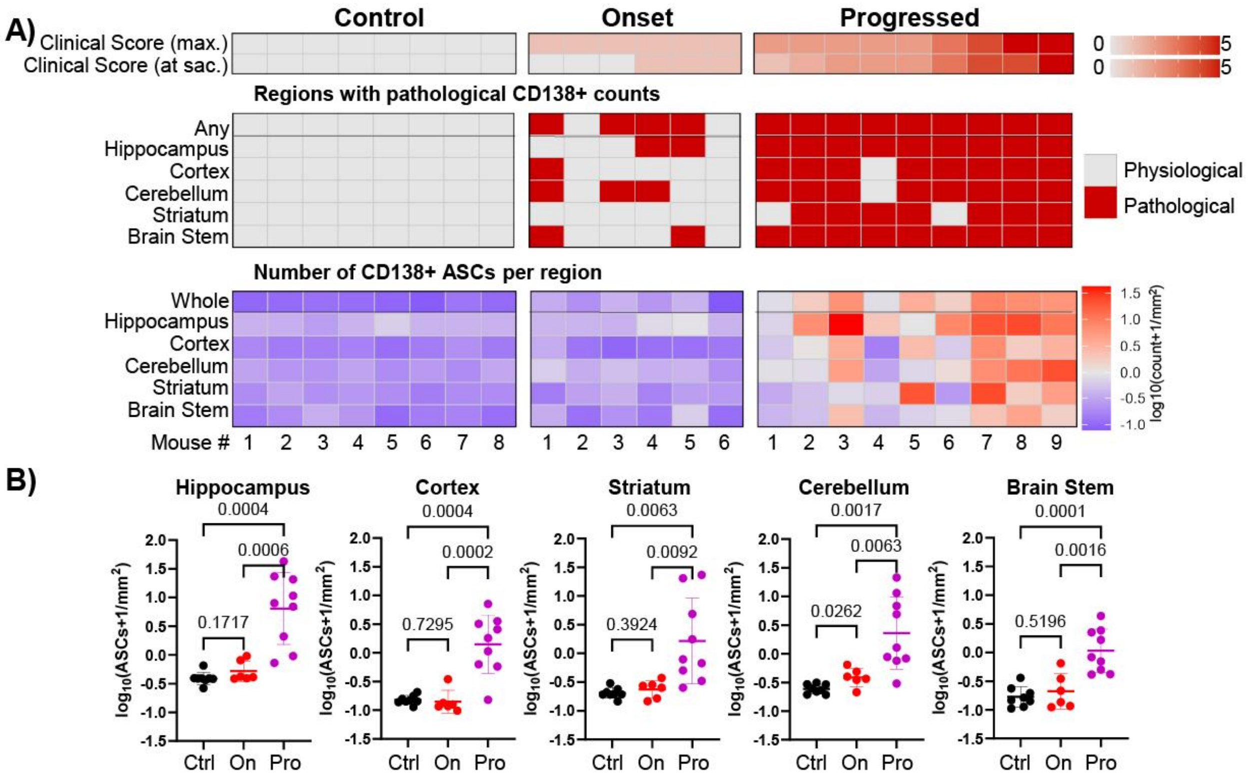


**Fig. S5. Anatomical localization of brain ASCs.** (**A**) Heatmap of pathological CD138$^{+}$ ASC counts across brain regions. ASC counts were classified as pathological (highlighted in red) if they were above the physiological range defined by the average ± 3*SD of Control mice. (**B**) Number of ASCs per area of brain region across brain regions. Data presented individually for each mouse and as mean ± SD. Welch's corrected t-tests.

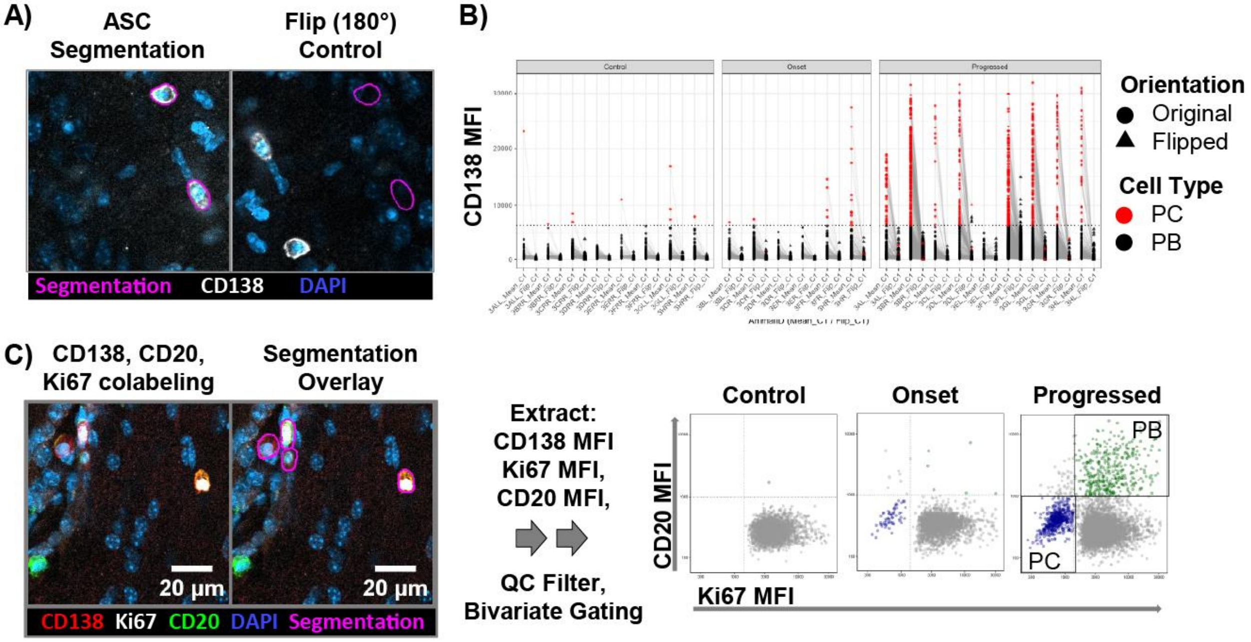


**Fig. S6. Pipeline for quantifying and phenotype brain ASCs from immunofluorescence.** (**A**) Illustration images of CD138$^{+}$ ASC segmentations and the flip control (segmentation overlaid on 180° rotated images) used for quality control purposes. (**B**) Quality control of ASC segmentation by comparing the segmented cells (original orientation) CD138 mean fluorescence intensities (MFI) to background fluorescence sampled with flip controls (180° rotated images). (**C**) Overview of the computational ASC segmentation pipeline. Stardist-based ASC segmentations from the CD138 single channel images are overlaid on the multi-channel image (i.e. CD138, CD20, Ki67, DAPI) to extract MFIs. Results are filtered using a uniform minimum cell size and minimum CD138 MFI. Filtered ASC subsets are then classified as CD20$^{+}$Ki67$^{+}$ plasmablasts (PBs) and CD20$^{-}$Ki67$^{-}$ plasma cells (PCs) by bivariate gating.

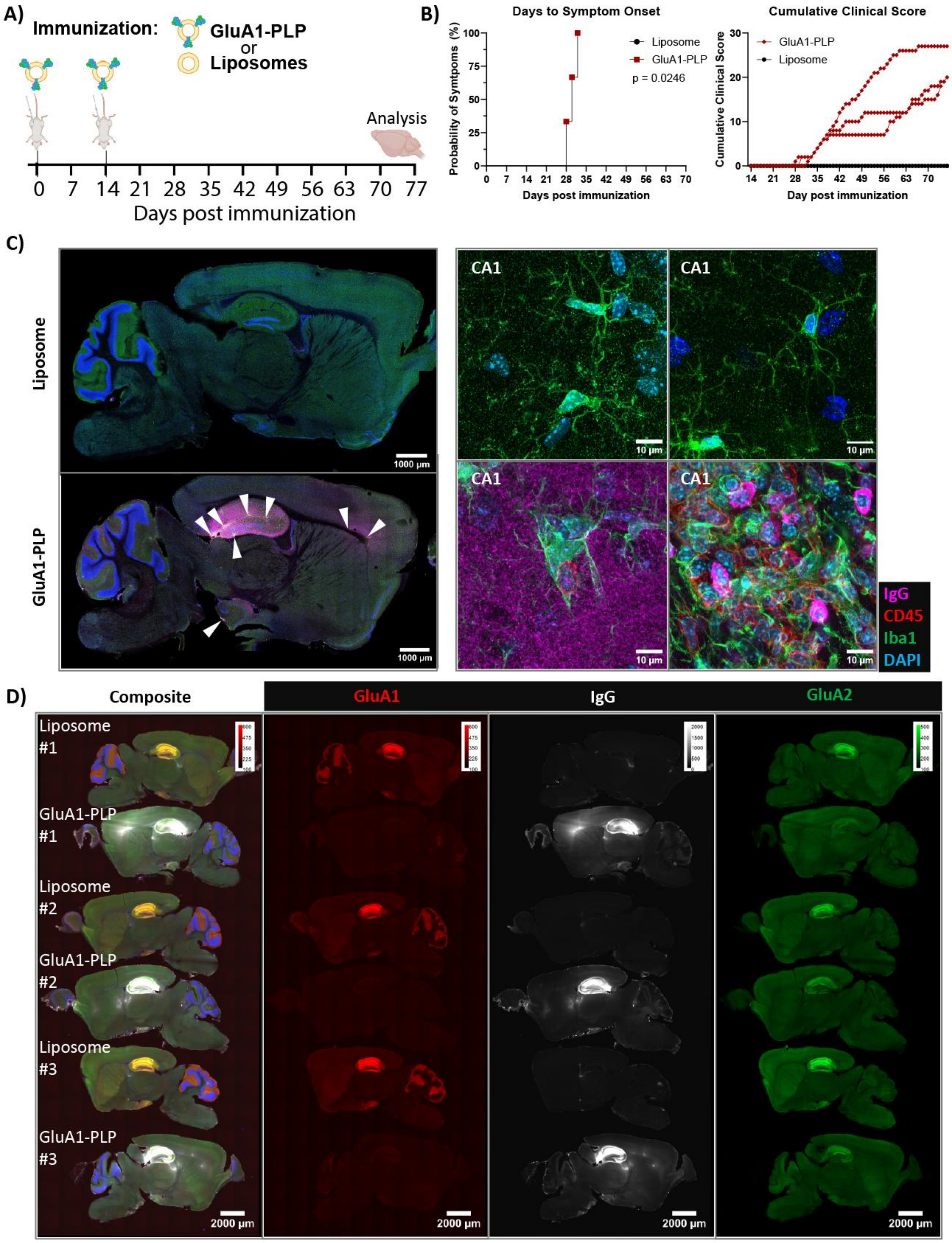

A)
Immunization:
GluA1-PLP
or
Liposomes
Analysis
0 7 14 21 28 35 42 49 56 63 70 77
Days post immunization
B)
Days to Symptom Onset
Probability of Symtpoms (%)
Liposome
GluA1-PLP
p = 0.0246
Days post immunization
Cumulative Clinical Score
GluA1-PLP
Liposome
Cumulative Clinical Score
Day post immunization
C)
Liposome
GluA1-PLP
1000 µm
CA1
10 µm
IgG
CD45
Iba1
DAPI
D)
Composite
GluA1
IgG
GluA2
Liposome #1
GluA1-PLP #1
Liposome #2
GluA1-PLP #2
Liposome #3
GluA1-PLP #3
2000 µm

**Fig. S7. GluA1-PLP immunization of mice induces AMPAR-AE with anti-GluA1 autoantibodies.** (**A**) Experimental setup (created with BioRender) of GluA1-proteoliposome (GluA1-PLP, n=3) or liposome immunization (n=3) of BALB/c mice and analysis of pathology. (**B**) Clinical disease course of GluA1-PLP induced AMPAR-AE. (**C**) Representative immunofluorescence of IgG, microglia ($Iba^{+}$), and lymphocytes ($CD45^{+}$) in brains of GluA1-PLP immunized mice (n=3 mice/group) demonstrating the presence of inflammatory microglia, IgG deposition, and lymphocyte clusters (white arrows) in the hippocampus and other brain regions in GluA1 immunized mice. (**D**) GluA1 immunoreactivity (4H9 Fab), IgG deposition (anti-mouse IgG Fc) and GluA2 immunoreactivity (15F1 Fab) in the brains of GluA1-PLP and liposome immunized mice (DAPI in blue). Each brain slice represents a different mouse (n=3 per group).

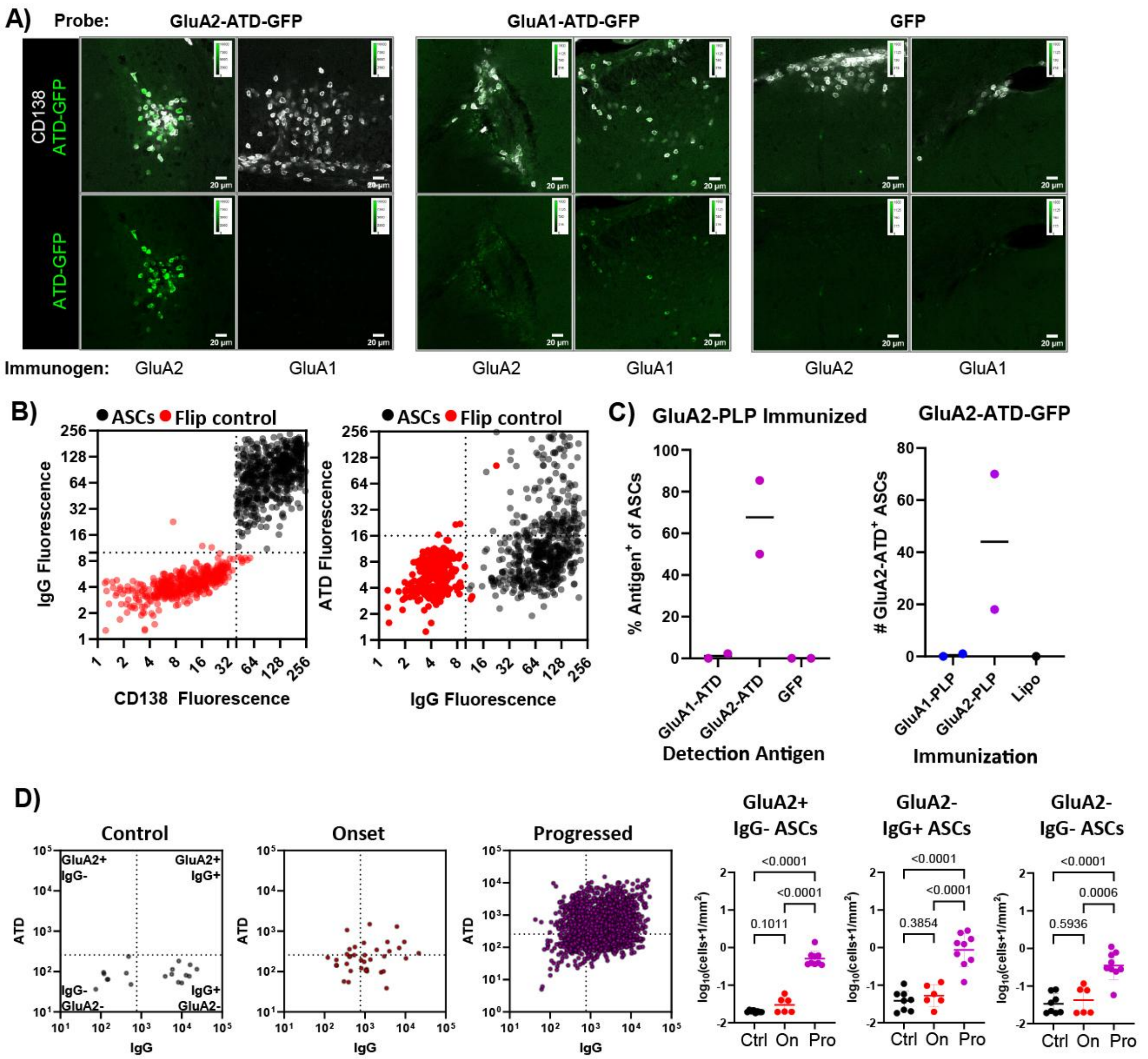

**Fig. S8. ATD-GFP probe detects ATD-specific ASCs *in situ*.** (**A**) Example images of histological staining with CD138 and GluA2-ATD-GFP, GluA1-ATD-GFP or a GFP control. Brain sections were obtained from GluA2-PLP and GluA1-PLP immunized mice as indicated by immunogen. (**B**) Quality control for segmentation and classification of ASCs (CD138$^{+}$IgG$^{+}$) as opposed to flipped controls by bivariate gating. Dotted lines indicate positivity thresholds. (**C**) Proportion and number of brain ASCs binding ATD-GFP probes quantified with the *in-situ* phenotyping pipeline in brain sections of GluA1-PLP, GluA2-PLP, and liposome (Lipo) immunized mice. (**D**) Quantification of GluA2-ATD and IgG positive and negative ASCs in Control (Ctrl), Onset (On), and Progressed (Pro) mice. Data presented as mean ± SD. Welch's t-tests.

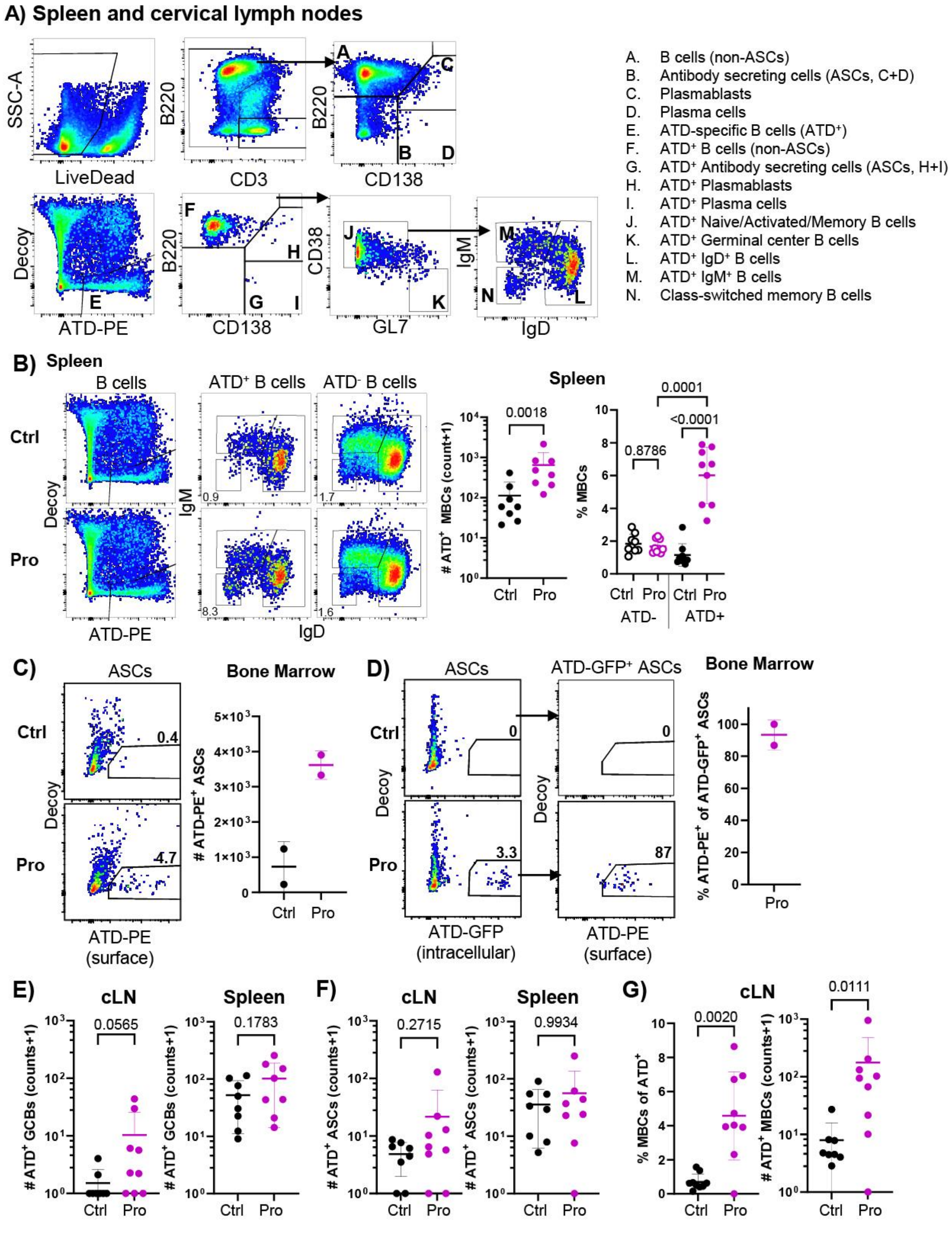

A) Spleen and cervical lymph nodes
A. B cells (non-ASCs)
B. Antibody secreting cells (ASCs, C+D)
C. Plasmablasts
D. Plasma cells
E. ATD-specific B cells (ATD+)
F. ATD+ B cells (non-ASCs)
G. ATD+ Antibody secreting cells (ASCs, H+I)
H. ATD+ Plasmablasts
I. ATD+ Plasma cells
J. ATD+ Naive/Activated/Memory B cells
K. ATD+ Germinal center B cells
L. ATD+ IgD+ B cells
M. ATD+ IgM+ B cells
N. Class-switched memory B cells
B) Spleen
B cells
ATD+ B cells
ATD- B cells
Ctrl
Pro
Spleen
0.0018
### ATD+ MBCs (count+1)
% MBCs
0.0001
<0.0001
0.8786
ATD-
ATD+
C) ASCs
Bone Marrow
ATD-PE (surface)
### ATD-PE+ ASCs
D) ASCs
ATD-GFP+ ASCs
ATD-GFP (intracellular)
ATD-PE (surface)
% ATD-PE+ of ATD-GFP+ ASCs
E) cLN
Spleen
0.0565
0.1783
### ATD+ GCBs (counts+1)
F) cLN
Spleen
0.2715
0.9934
### ATD+ ASCs (counts+1)
G) cLN
0.0020
0.0111
% MBCs of ATD+
### ATD+ MBCs (counts+1)

**Fig. S9. GluA2-ATD-specific B cell response in secondary lymphoid tissues.** (**A**) Flow cytometry gating strategy for spleen and cervical lymph node (cLN) B cells (spleen shown, pre-gated on single cell lymphocytes) with subsets indicated. ATD-specific B cells pre-gated on total B cells (A+B). (**B**) Representative flow cytometry of splenic B cells gated for ATD$^+$ B cells (GluA2-ATD-PE$^+$Decoy$^-$) or non-probe binding/decoy-binding (ATD$^-$) B cells and then MBCs (class-switched) in Control and Progressed mice. Number of ATD$^+$ MBCs and proportion of ATD$^+$ or ATD$^-$ B cells that are MBCs across groups. (**C** and **D**) Representative flow cytometry of GluA2-ATD-PE surface and GluA2-ATD-GFP intracellular stains on bone marrow ASCs (CD138$^+$). (**C**) Number of ATD-PE$^+$ ASCs in Control and Progressed mice. (**D**) Proportion of ATD-GFP$^+$ bone marrow ASCs that bound ATD-PE on their surface in Progressed mice (data in C and D from two experiments). (**E**) Number of ATD$^+$ germinal center B cells (GCBs), (**F**) number of ATD$^+$ ASCs and (**G**) Percentage and number of ATD$^+$ B cells that are class-switched MBCs in the cLNs and spleens of Control (Ctrl) and Progressed (Pro) GluA2-PLP mice as indicated. Data from each mouse presented individually and as mean ± SD. (B-left, E, F, G-right) lognormal Welch's t-test, (B-right) Welch's ANOVA with Dunnett's T3 post-hoc t-test, (G-left) Welch's t-test.

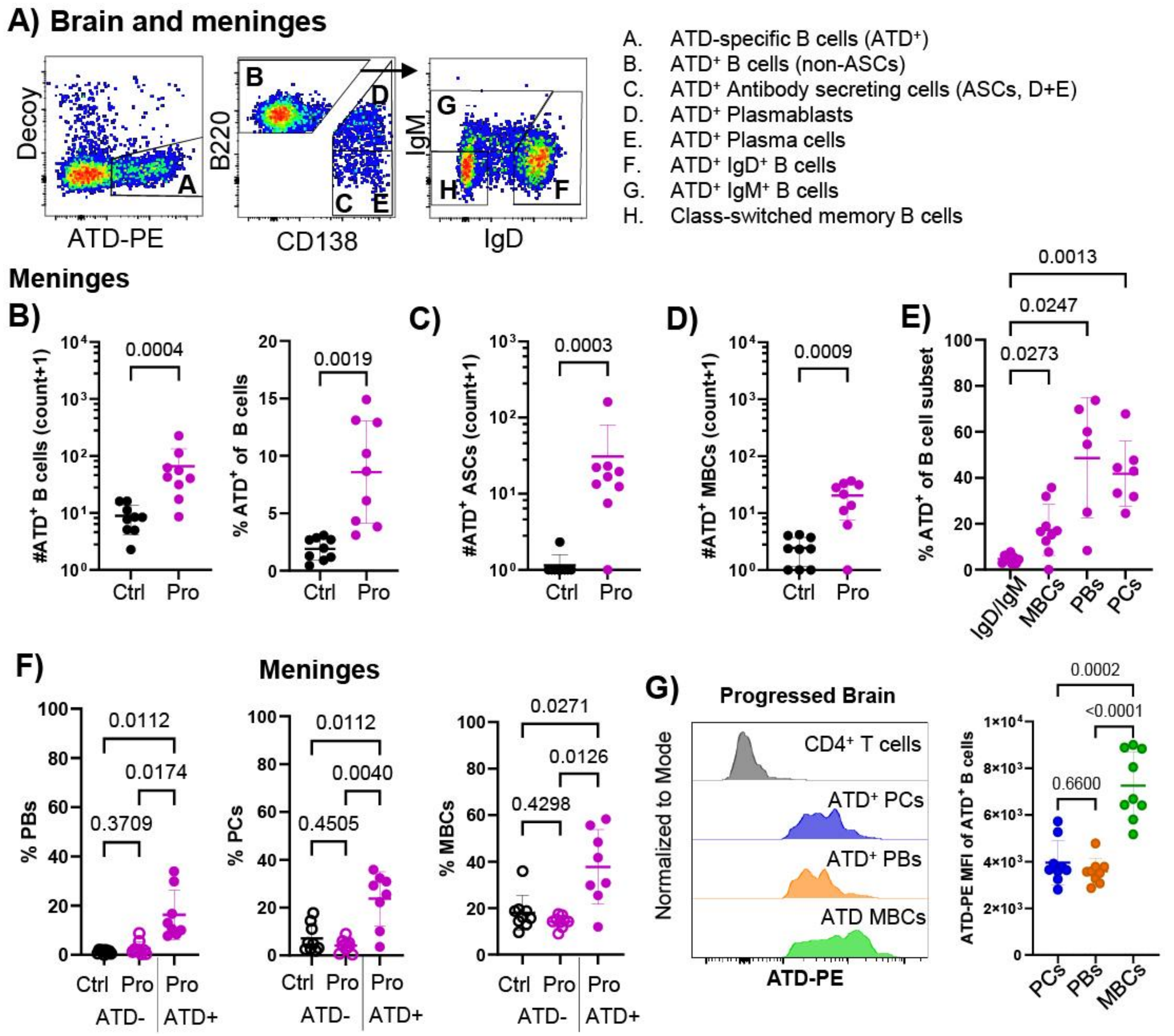


**Fig. S10. GluA2-ATD-specific B cell response in CNS tissues.** (**A**) Flow cytometry gating strategy for ATD-specific B cells in brain and meninges (brain shown, pre-gated on i.v. label- B cells as in Fig. S2). (**B**) Number and proportion of total B cells that are ATD$^+$, (**C**) number of ATD$^+$ ASCs and (**D**) number of ATD$^+$ class-switched memory B cells (MBCs) in the meninges. (**E**) Proportion of B cell subsets that are ATD-specific in the meninges. (**F**) Percentage of ATD$^+$ and ATD$^-$ B cells that are plasmablasts (PBs), plasma cells (PCs), and MBCs in the brain. (**G**) Representative ATD binding by total CD4+ T cells and ATD$^+$ PCs, PBs and MBCs in the brain of a Progressed mouse and comparison of ATD-PE MFI between ATD$^+$ B cell populations. Data from each mouse presented individually and as mean ± SD. Ctrl, Control;

Pro, Progressed. (B-right, C, D) lognormal Welch's t-test, (B-left) Welch's t-test, (E, F, G) Welch's ANOVA with Dunnett's T3 post-hoc t-test.
